# Influenza A virus NS1 sequesters RNA:DNA hybrids to evade RNase H1-dependent innate immunity

**DOI:** 10.64898/2026.05.04.722684

**Authors:** Florence Kwaschik, Ian Pichler, José L. Ruiz, Verena Kufner, Michael Huber, Benjamin G. Hale

**Affiliations:** Institute of Medical Virology, University of Zurich, Zurich 8057, Switzerland; Functional Genomics Center Zurich, ETH Zurich and University of Zurich, Zurich 8057, Switzerland; Swiss Institute of Bioinformatics, Amphipôle, Quartier UNIL-Sorge, Lausanne 1015, Switzerland

## Abstract

RNA:DNA hybrids, and their aberrant accumulation, are key regulators of human genome integrity and innate immunity. Given the functional parallels and extensive interplay between the human genome and viral genetic material, RNA:DNA hybrids also have established roles during infections with DNA viruses and retroviruses. Nevertheless, the presence and consequences of RNA:DNA hybrids in RNA virus infections, which lack a DNA phase, remain largely uncharacterized. Here, we show that infection of human cells with influenza A virus (IAV), but not several other respiratory negative-sense RNA viruses, induces the unexpected accumulation of RNA:DNA hybrids in perinuclear regions. Sequencing of infection-induced cytoplasmic RNA:DNA hybrids revealed that they derive predominantly from the human host genome, with apparent preference for intronic and intergenic sequences, although an intriguing fraction also originates from the IAV genome, with some enrichment for specific viral segments. Notably, we identify the IAV non-structural protein, NS1, as a key determinant of RNA:DNA hybrid localization and stability: NS1 is required for, and co-localizes with, perinuclear RNA:DNA hybrids, and its absence appears to sensitize RNA:DNA hybrids to metabolism by the cellular ribonuclease, RNase H1. Furthermore, experimental loss of RNase H1 attenuates IAV-induced antiviral responses, including type I interferon and inflammatory gene expression programs, indicating that RNA:DNA hybrid metabolism likely contributes to host defense. Overall, our findings uncover host- and viral-origin RNA:DNA hybrids as a previously unrecognized feature of a human pathogenic RNA virus infection, and suggest a mechanism by which a viral product antagonizes host responses mediated via these unusual nucleic acids by spatial sequestration.

## Introduction

To mount an effective defense against viruses, infected cells must efficiently detect pathogens and initiate an antiviral response. This process relies on specialized host sensor proteins that engage foreign ‘non-self’ structures, typically nucleic acids, which are recognized due to distinctive features or localizations not commonly found with ‘self’ nucleic acids (1). A central challenge in this mechanism is the precise discrimination of self from non-self, given that many viral infections perturb host nucleic acid homeostasis (2–4). Specialized pattern recognition receptors (PRRs), such as RIG-I, MDA5, cGAS, ZBP1 or TLRs, are key sensor proteins that recognize characteristic traits of specific viral nucleic acids and, once activated, initiate signaling cascades that result in the production of antiviral cytokines, such as interferons (IFNs) or inflammatory mediators (5). Importantly, such responses require tight regulation in order to prevent potential immunopathological consequences, thus ensuring a balanced and controlled host defense mechanism (6). While PRRs primarily detect pathogen-associated molecular pattern (PAMP) nucleic acids, emerging evidence indicates that, under certain conditions, these sensors can also recognize self nucleic acids (7–10). This occurs when endogenous nucleic acids are mislocalized, structurally altered, aberrantly metabolized, or transcriptionally upregulated during pathological states, potentially contributing to inflammatory responses (11). In some of these contexts, the inability to correctly discriminate between self and non-self can lead to autoinflammatory responses, including interferonopathies such as systemic lupus erythematosus (SLE) and Aicardi-Goutières syndrome (AGS), in which genes involved in correct nucleic acid metabolism or immune system regulation are defective (12). In other contexts, virus infections can blur the distinction between self and non-self, and sensing of endogenous molecules can be a mechanism by which the cell amplifies immunostimulation, with virus-triggered transposable elements (TEs), mitochondrial DNA release, and transcription-termination by-products having all previously been shown to contribute to innate antiviral defenses (3, 13–16).

During viral infections, diverse nucleic acid species (including double-stranded RNA (dsRNA), single-stranded RNA (ssRNA), dsDNA, ssDNA, and RNA:DNA hybrids) have been observed, many of which have been reported to act as effective PAMPs (17). However, the origin and functional relevance of these nucleic acid species can differ markedly depending upon the viral genome type and replication strategy. In the context of influenza A virus (IAV), a negative-sense single-stranded RNA virus that replicates within the cell nucleus, both virus- and host-derived dsRNAs have been detected experimentally with immunostaining or sequencing approaches (13, 18, 19). These nucleic acid species have variously been proposed to serve as signals that trigger the host innate immune system through specific PRRs (13, 18, 19). Viral dsRNAs may originate from aberrantly generated replication intermediates, annealing of complementary vRNA and cRNA/mRNA species, or from the partially double-stranded pan-handle structures of the IAV genome (20). Infection-triggered host dsRNAs can derive from upregulated TE RNAs that might form dsRNAs through bidirectional transcription, intramolecular base-pairing, or overlapping sense and anti-sense transcription (21). Intriguingly, some TEs encode reverse transcriptases, allowing the synthesis of complementary DNA from RNA, and thus the formation of complementary RNA:DNA hybrids. This raises the possibility that an RNA virus infection, such as with IAV that lacks a DNA stage, could indirectly give rise to RNA:DNA hybrid nucleic acids through host-encoded enzymatic activities. Indeed, RNA:DNA hybrids have been reported to be generated by such a process during experimental infection of induced pluripotent stem cells with a picornavirus, in which the viral RNA was reverse transcribed to DNA by a cellular reverse transcriptase allowing formation of an RNA:DNA hybrid (22). Notably, host-derived reverse transcriptase activity has also been documented to occur during some arenavirus, rhabdovirus, coronavirus, and paramyxovirus infections (23–25). In addition, RNA:DNA hybrids can arise in cells during normal gene regulation and DNA repair processes, most notably via co-transcriptional R-loop formation by RNA polymerase II (26). Furthermore, aberrantly elevated or persistent levels of RNA:DNA hybrids can result from replication-transcription collisions and excessive DNA damage-associated processes, and their accumulation can exacerbate genomic instability as well as trigger innate immune responses (27, 28). Consistent with this, dysregulated RNA:DNA hybrid metabolism has been linked to activation of cGAS, TLR3, TLR9 and ZBP1-mediated signaling responses in multiple contexts (27, 29, 30). Nevertheless, the generation, origins, and potential antiviral contributions of RNA:DNA hybrids during RNA virus infections have yet to be fully explored.

Given the intimate association between IAV replication and host nuclear processes, not least RNA polymerase II-dependent transcription and de-repression of TEs (13, 18, 31), we sought to determine the presence and role of RNA:DNA hybrids during IAV infection. Using a combination of immunofluorescence microscopy, sequencing analysis, and genetic perturbation approaches, we demonstrate that host- and virus-derived RNA:DNA hybrids accumulate in perinuclear regions of IAV-infected human cells. Furthermore, we provide evidence that the viral NS1 protein aids the spatial sequestration of RNA:DNA hybrids and limits their metabolism by RNase H1, a cellular ribonuclease that potentiates immune and inflammatory gene expression during IAV infection. These data reveal an unexpected facet of host-pathogen biology, whereby RNA virus-induced RNA:DNA hybrid accumulation and its antagonism by a viral protein modulate host innate immunity.

## Results

### IAV infection leads to perinuclear accumulation of RNA:DNA hybrids

To characterize nucleic acid species formed during respiratory RNA virus infections, we used immunofluorescence microscopy to screen for the presence of dsRNA and RNA:DNA hybrids in human lung epithelial cells infected with a panel of pathogenic viruses. Specifically, A549 cells were infected with influenza A virus (IAV), respiratory syncytial virus (RSV), parainfluenza virus type 2 (PIV2), parainfluenza virus type 5 (PIV5), or measles virus (MeV), and were subsequently fixed with methanol at timepoints corresponding to robust infection levels for each virus. Following proteinase K treatment and immunostaining with antibodies specific for dsRNA (9D5, (32)) and RNA:DNA hybrids (S9.6, (33)), we visualized cells by fluorescence confocal microscopy. In parallel samples, infection levels were determined by paraformaldehyde-based fixation and immunostaining for IAV nucleoprotein (NP) or GFP (for GFP-expressing RSV, PIV2, PIV5 and MeV). Under these conditions, we could consistently detect dsRNA in the cytoplasm of cells from cultures infected with IAV and MeV, but not with RSV, PIV2 or PIV5 (**Figs. 1A-B**). Strikingly, we observed a clear perinuclear signal for RNA:DNA hybrids that was specific to IAV infected cultures, and which appeared to co-localize with the dsRNA signal (**Figs. 1A, 1C, S1A**). Given the potential for the S9.6 antibody to sometimes exhibit non-specific binding to dsRNA (34), we further assessed specificity of the detectable RNA:DNA hybrid signal by similarly immunostaining A549 cells infected with Semliki Forest virus (SFV) or herpes simplex virus type 1 (HSV-1), two viruses with divergent genome types that have been reported to produce abundant levels of dsRNA during infection (35, 36). In cultures infected with SFV or HSV-1, cytoplasmic dsRNA was indeed observable, but no signal was detected with the S9.6 antibody (**Figs. 1D-F**). The absence of cross-reactivity of the S9.6 antibody to high levels of SFV dsRNA in our experimental system suggests that the detection of RNA:DNA hybrids in IAV-infected cultures is specific. By co-staining IAV-infected A549 cells for dsRNA or RNA:DNA hybrids together with NP, we noted that perinuclear accumulation of dsRNA and RNA:DNA hybrids occurred only in individual infected cells, and was absent from non-infected bystander cells (**Figs. 1G-H**). Furthermore, beyond A549 cells, we observed cytoplasmic RNA:DNA hybrids and dsRNA in IAV-infected MRC5 (primary-like lung fibroblast), Calu-3 (lung epithelial), and HT-29 (colorectal epithelial) cells (**Figs. S1B-D**). In addition, perinuclear RNA:DNA hybrids and dsRNA were detectable following infection of A549 cells with various different human and avian IAV strains, although it was notable that the relative abundance and localization of RNA:DNA hybrids and dsRNA detected varied with each strain (**Figs. S1E-G**). These observations reveal the disparate production of observable dsRNA species during infection of human cells with several different viruses, and reveal an intriguing specificity of perinuclear RNA:DNA hybrid accumulation during infections with IAV.

**Figure 1:**
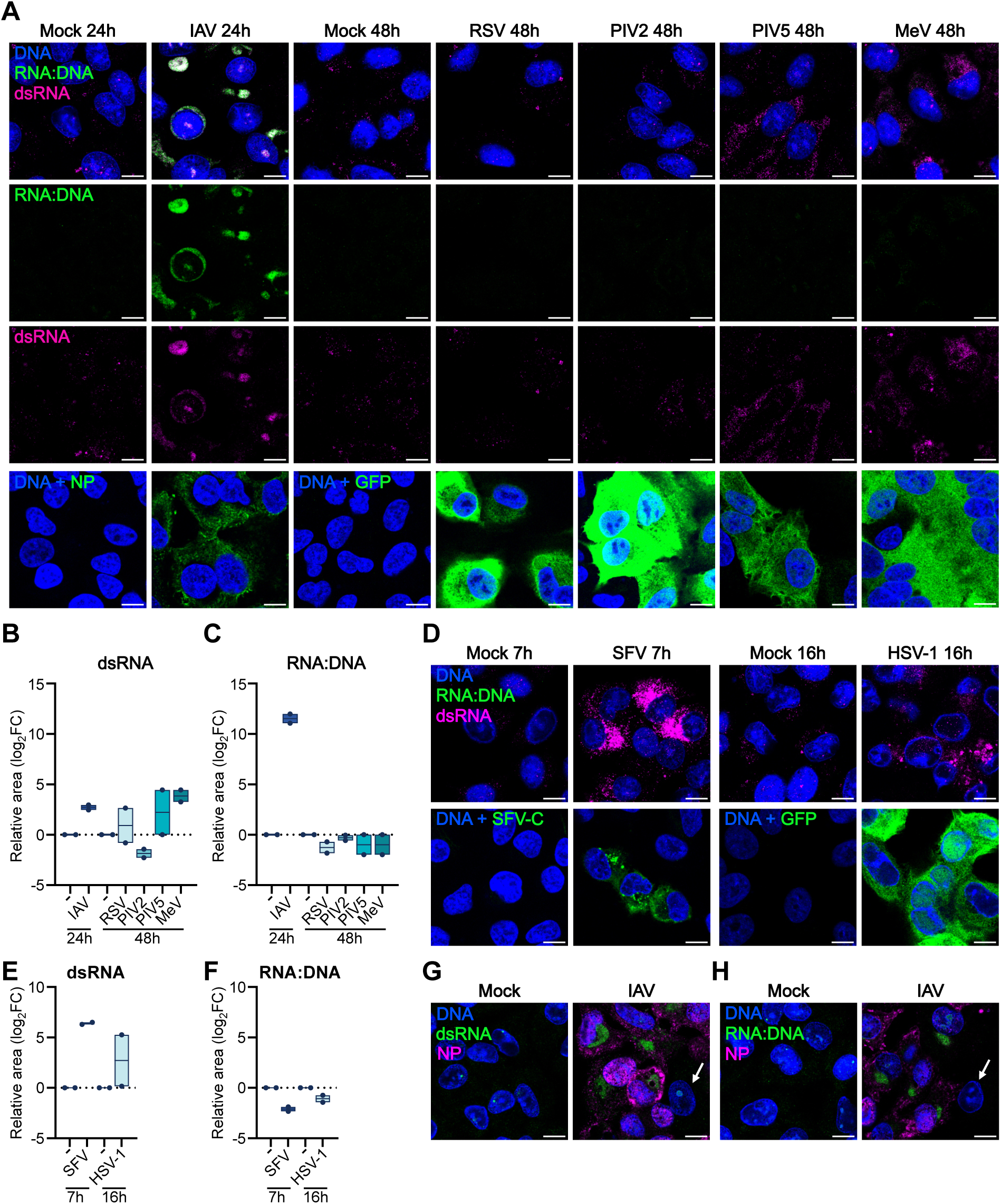
IAV infection leads to perinuclear accumulation of RNA:DNA hybrids. **A**. Immunofluorescence confocal microscopy of A549 cells infected with IAV, RSV-GFP, PIV2-GFP, PIV5-GFP (all at MOI 1 PFU/cell), MeV-GFP (MOI 0.5 PFU/cell), or mock for the indicated times. Upper three rows: cells were fixed and permeabilized with methanol, proteinase K treated, and then stained for RNA:DNA hybrids (S9.6; green), dsRNA (9D5; magenta), and DNA (DAPI; blue). Lower row: cells were fixed with 4% PFA, permeabilized with confocal buffer, and then stained for IAV NP (green) or GFP (green) as appropriate. Nuclei were stained with DAPI (blue). **B-C**. Quantification of cytoplasmic dsRNA (B) and RNA:DNA hybrid (C) signals from experiments shown in A. Boxplots indicate the range and mean log_2_ fold change (FC) of the area fraction occupied by signal relative to mock-infected controls from n=2 independent experiments. Each dot represents the average log_2_FC from multiple fields imaged per well. The dotted line indicates no change relative to mock. For replicates where mock or infected signal was not detected, FC was calculated relative to the limit of detection (LoD). **D**. Immunofluorescence confocal microscopy of A549 cells infected with SFV (MOI: 1 PFU/cell; 7 h), HSV-1-GFP (MOI: 1 PFU/cell; 16 h), or mock. Procedures for fixation, enzyme treatment and staining were performed as described in A. Upper row shows staining for RNA:DNA hybrids (S9.6; green), dsRNA (9D5; magenta), and DNA (DAPI; blue). Lower row shows staining for the SFV C protein (green) or GFP (green). **E-F**. Quantification of cytoplasmic dsRNA (E) and RNA:DNA hybrid (F) signals from experiments shown in D, with methods as described in B-C. **G-H**. Immunofluorescence confocal microscopy of A549 cells infected with IAV, or mock, as described in A. Immunostaining was performed for dsRNA (9D5; green) and NP (magenta) (G), or RNA:DNA hybrids (S9.6; green) and NP (magenta) (H). Nuclei were stained with DAPI (blue). White arrows depict uninfected cells. For all panels, data are representative of at least n=2 independent experiments. For microscopy images, scale bars represent 10 µm.

### IAV infection-triggered RNA:DNA hybrids originate from host and viral sequences

While IAV-associated dsRNAs likely originate from viral genome replication via RNA intermediates or infection-triggered upregulation of TE RNAs (13, 18, 19, 37), the origin of RNA:DNA hybrids detected during IAV infection is unclear. We therefore employed cytoplasmic DNA:RNA hybrid immunoprecipitation sequencing (cytoDRIP-seq) as an approach to investigate RNA:DNA hybrid identity (27). A549 cells were infected with IAV (or mock) for 24 h, and the cytoplasmic fractions were isolated (**Fig. 2A**). For verification of subcellular fractions, RT-qPCR was performed for the established nuclear marker, MALAT1 (38), as well as the cytosolic marker, GAPDH (39) (**Fig. S2A**). Following total nucleic acid extraction, samples were subjected to immunoprecipitation with the S9.6 antibody to enrich for RNA:DNA hybrids, or with a V5 antibody as a negative control. Prior to immunoprecipitation, split samples had been digested with RNase H1 (or mock) to degrade the RNA strand of hybrids in one sample, thus providing a further specificity control for later sequencing steps. Following nucleic acid extraction from the immunoprecipitated samples, DNA libraries were prepared and sequenced using a single-end approach (40), providing sufficient resolution to determine genomic distributions of identified sequences. Using the split sample approach with and without RNase H1 treatment, alongside the antibody specificities, we were able to define a background-subtracted signal for RNA:DNA hybrids (ΔDRIP-seq), which was used for subsequent analyses. To assess the efficiency and composition of the captured libraries, we initially analyzed the total and relative mapped read counts for non-RNase H1-treated samples. Immunoprecipitation with the S9.6 antibody from infected cell lysates yielded libraries with around 10^6^ human-mapped reads, representing an approximately 100-fold increase in absolute yield over both isotype and uninfected controls (approx. 10^4^ reads) (**Fig. 2B**). Despite this difference in scale, human-derived sequences accounted for an average of 87-99% of the signal associated with RNA:DNA hybrids across all samples (**Fig. S2B**). Within these libraries, reads mapping to the reference IAV genome were also detected at an enriched count and frequency in infected samples immunoprecipitated with the S9.6 antibody (approx. 2x10^3^ viral reads; 0.3%) as compared with infected samples immunoprecipitated with the isotype control antibody (approx. 10 viral reads; 0.08%) (**Figs. 2C, S2C**). These specific enrichments of human and viral sequences in S9.6 antibody immunoprecipitates from infected samples were maintained even after correcting for non-specific background signal by integrating data from the RNase H1-treated samples (**Figs. S2D-E**). Further analysis of reads mapping to the human genome revealed that there was an apparent enrichment of RNA:DNA hybrid sequences originating from intronic and intergenic regions following IAV infection and S9.6 immunoprecipitation, as compared with the pattern observed with the non-specific isotype control (**Figs. 2D-E**). Of the small proportion of identified RNA:DNA hybrid sequences mapping to the IAV genome, the sequences originated from all IAV segments. Nevertheless, the sequences identified were not uniformly distributed, and there was apparent slight preferential mapping to specific (mostly 5’) regions of IAV genomic segments 4, 7 and 8, even when accounting for relative segment length (**Figs. 2F, S2F**). These data reveal that the RNA:DNA hybrids detected in the cytoplasm of IAV-infected cells originate from host and viral sequences.

**Figure 2:**
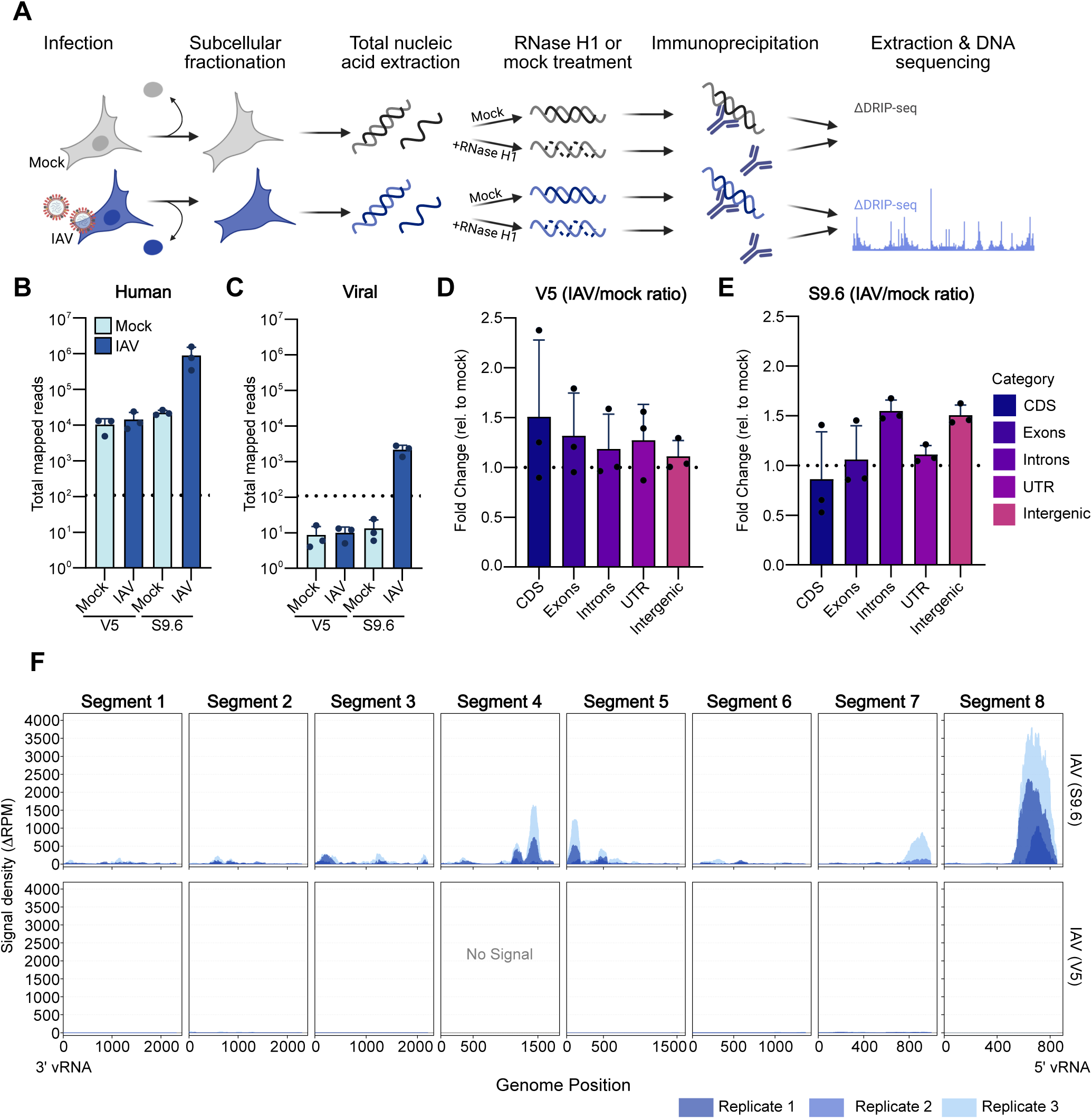
IAV infection-triggered RNA:DNA hybrids originate from host and viral sequences. **A**. Schematic representation of the modified cytoDRIP-seq approach. A549 cells were infected with IAV at an MOI of 1 PFU/cell for 24h. Immunoprecipitations were performed with anti-V5 (control) or S9.6 antibodies. ΔDRIP-seq represents RNase H1-sensitive RNA:DNA hybrid signal calculated as non-RNase H1-treated samples minus RNase H1-treated samples. Three independent experiments were performed and analyzed. B-C. Total number of reads mapped to the (B) human genome and the (C) IAV genome. Data represent the composition of non-RNase H1-treated samples to illustrate the initial distribution of captured sequences. The dashed horizontal line indicates the minimum threshold for high-confidence signal detection (109 reads); data points below this line represent background noise or low-complexity libraries. Bars represent the mean of n=3 independent biological replicates, with individual dots indicating the values for each replicate. Error bars represent standard deviations. **D-E**. Bar plots showing the average distribution of normalized signal across human genomic features including coding sequence (CDS), exons, introns, untranslated regions (UTR), and intergenic regions. Data are represented as the mean fold change in signal density relative to the mock condition for (D) V5- and (E) S9.6-immunoprecipitated samples. Dots represent n=3 biological replicates derived from background-subtracted (ΔDRIP-seq) signal. Error bars represent standard deviations. **F**. Genomic distribution of viral-origin RNA:DNA hybrid sequences mapping to each of the eight IAV genome segments. Coverage plots show the ΔRPM signals (RPM difference between non-RNase H1-treated and RNase H1-treated samples; negative values set to zero) obtained from IAV-infected cells following immunoprecipitation with either S9.6 or V5 (control) antibodies. Overlaid traces in varying shades represent each of the n=3 independent replicates. Genome position is indicated on the x-axis.

### IAV NS1 contributes to perinuclear presence of infection-induced RNA:DNA hybrids

We previously observed that the IAV NS1 protein, a key viral host defense antagonist (41), limits cytosolic accumulation of dsRNA in infected cells by a mechanism that appears to involve physical sequestration of dsRNA in perinuclear regions (18). Given that these regions are similar to those where RNA:DNA hybrids accumulate during IAV infection, we next assessed the interplay between NS1 and RNA:DNA hybrids. Akin to its co-localization with perinuclear dsRNA, immunofluorescence microscopy revealed that NS1 also partially co-localized with perinuclear RNA:DNA hybrids (**Figs. 3A-D, S3A-B**). Strong co-localization was specific to the NS1 protein, as the IAV PB2 protein exhibited a mostly distinct localization pattern under the same experimental conditions (**Figs. 3E-H, S3C-D**). Consistent with the co-localization of NS1 with both dsRNA and RNA:DNA hybrids, both the dsRNA antibody, 9D5, and the RNA:DNA hybrid antibody, S9.6, could co-precipitate NS1 from IAV-infected cells (**Fig. 3I**). Intriguingly, while infection of A549 cells with an IAV strain genetically-engineered to lack expression of NS1 (IAV ΔNS1) led to induced dsRNAs distributing diffusely throughout the cytoplasm compared to the perinuclear concentration observed with wild-type (wt) IAV (18), RNA:DNA hybrids could not be detected at all in IAV ΔNS1-infected cells (**Figs. 3J-K**). As a large portion of the NS1 genomic sequence is lacking in IAV ΔNS1, this observation could implicate this specific sequence in the RNA:DNA hybrid population, which would be supported by our ΔDRIP-seq results (**Fig. 2F**). However, given the large fraction of IAV-induced RNA:DNA hybrids that also derive from host sequences, it seems unlikely that lack of this NS1 genomic sequence alone would have such a complete phenotype. Similar to the suggestion that the NS1 protein physically sequesters dsRNAs into perinuclear regions (18), the observation that NS1 also co-localizes and engages with RNA:DNA hybrids may indicate that NS1 shields these hybrids from factors that would otherwise cause RNA:DNA hybrid destabilization, a hypothesis explored below.

**Figure 3:**
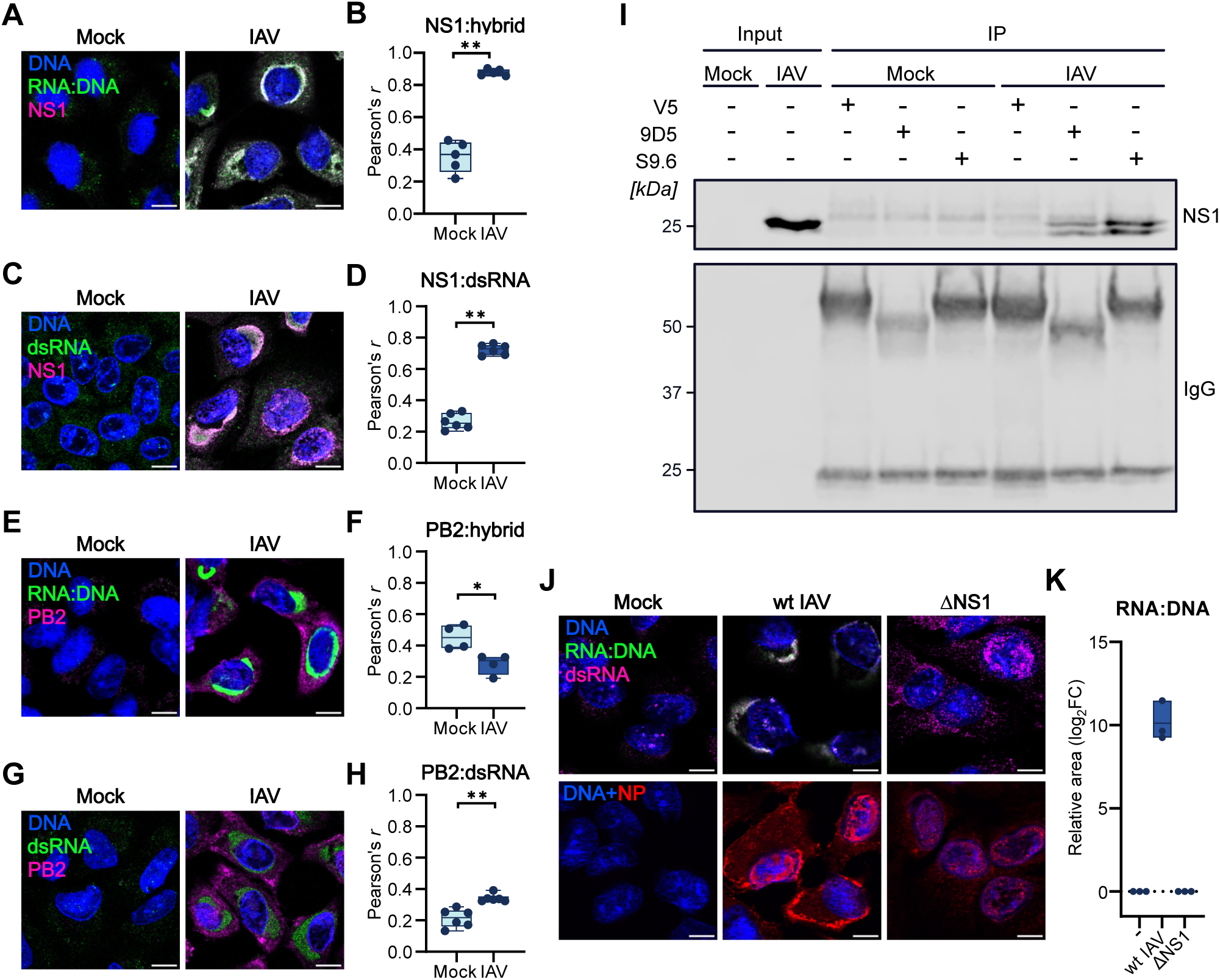
IAV NS1 contributes to perinuclear presence of infection-induced RNA:DNA hybrids. **A-B**. Immunofluorescence microscopy analysis (A) and associated colocalization analysis (B) of A549 cells infected, or mock, with IAV at an MOI of 1 PFU/cell for 24 h, followed by methanol fixation, proteinase K treatment, and staining for RNA:DNA hybrids (S9.6; green), NS1 (magenta), and DNA (DAPI; blue). Colocalization between NS1 and RNA:DNA hybrids was assessed by determining the Pearson’s correlation coefficient (*r*) between green and magenta signals. Boxplots indicate the minimum to maximum range (whiskers) and mean (horizontal line) *r* values from n=3 independent experiments. Each dot represents the *r* value from a single field of view. Non-merged images are presented in Supplementary Figure 3. **C-D**. Detection of dsRNA (9D5; green), NS1 (magenta), and DNA (DAPI; blue) (C), as well as corresponding colocalization analysis (D) in A549 cells treated as detailed in A-B. Non-merged images are presented in Supplementary Figure 3. **E-F**. Detection of RNA:DNA hybrids (S9.6; green), PB2 (magenta), and DNA (DAPI; blue) (E), as well as corresponding colocalization analysis (F) in A549 cells treated as detailed in A-B. Non-merged images are presented in Supplementary Figure 3. **G-H**. Detection of dsRNA (9D5; green), PB2 (magenta), and DNA (DAPI; blue) (G), as well as corresponding colocalization analysis (H) in A549 cells treated as detailed in A-B. Non-merged images are presented in Supplementary Figure 3. **I**. Immunoblot analysis following immunoprecipitation with specific antibodies. A549 cells were infected with IAV, or mock, at an MOI of 1 PFU/cell for 24 h followed by immunoprecipitation (IP) of resulting cell lysates with anti-V5, anti-dsRNA (9D5), or anti-RNA:DNA hybrid (S9.6) antibodies. Input cell lysate and IP fractions were analyzed by immunoblotting for NS1 and total IgG. **J**. Immunofluorescence microscopy analysis of A549 cells infected with wt IAV, IAV ΔNS1, or mock, at an MOI of 1 PFU/cell for 24 h, prior to processing as described in A. Upper row shows staining for RNA:DNA hybrids (S9.6; green), dsRNA (9D5; magenta), and DNA (DAPI; blue). Bottom row shows staining for IAV NP (red) and DNA (DAPI; blue). **K**. Quantification of cytoplasmic RNA:DNA hybrid signals (S9.6; green) from experiments described in J. Boxplots indicate the range and mean log_2_FC of the area fraction occupied by signal relative to mock-infected controls from n=3 independent experiments. Each dot represents the average log_2_FC from multiple fields imaged per well. Dotted line indicates no change relative to mock. For all panels, data are representative of at least n=3 independent experiments. For microscopy images, scale bars represent 10 µm. For panels B, D, F, and H, statistical significance was determined by Mann-Whitney U test (\**P□*≤□0.05; \*\**P□*≤□0.01).

### IAV NS1 protects RNA:DNA hybrids from the action of cellular RNase H1

RNase H1 and RNase H2, cellular ribonucleases that recognize RNA:DNA hybrids and degrade the RNA strand (42), can be activated in response to RNA:DNA hybrid accumulation stress and have reported regulatory roles in metabolizing hybrids to limit pathological innate immune activation or loss of genome integrity (43, 44). RNase H1 was particularly intriguing to us given its previous association with a cytoplasmic antiviral mechanism (22). Indeed, immunofluorescence microscopy of A549 cells infected with wt IAV revealed co-localization of RNase H1 with RNA:DNA hybrids in perinuclear accumulations, something not detectable in A549 cells infected with IAV ΔNS1 where RNase H1 levels appeared to be more diffusely distributed across the cytoplasm (**Figs. 4A-B**). To explore the potential functional interplay between IAV and RNase H1 further, we generated a pool of *RNASEH1* knockout A549 cells using lentivirus-delivered CRISPR/Cas9, hereafter referred to as sgRH1 cells. As a methodological negative control, a pool of A549 cells was also generated with guide RNA targeting GFP, termed sgGFP. Western blot analysis confirmed specific reduction of RNase H1 levels in sgRH1 cells as compared with sgGFP cells (**Fig. 4C**). The reduced levels of RNase H1 in sgRH1 cells did not limit wt IAV infection-triggered accumulation of either RNA:DNA hybrids or dsRNA (**Figs. 4D-F**). In addition, both nucleic acid species were detectable in perinuclear regions, suggesting that RNase H1 is not the primary driver of this localization phenotype and that RNase H1 is likely recruited secondarily (**Fig. 4D**). Remarkably, however, the absence of RNase H1 led to the striking accumulation of RNA:DNA hybrids during IAV ΔNS1 infection (**Figs. 4D-F**). In addition, the signal for these RNA:DNA hybrids was clearly not perinuclear in distribution, but appeared more punctate throughout the cytoplasm and was distinct from the diffuse cytoplasmic dsRNA signal (**Figs. 4D, S4A-B**). Altogether, these data suggest that the viral NS1 protein is likely the driver of RNA:DNA hybrid and dsRNA localization to perinuclear regions. While the NS1 protein may sequester dsRNAs to dampen their detection by cytosolic sensors (18), another function of NS1 may be to limit catalytic access of RNase H1 to the RNA:DNA hybrids, thus preventing their metabolism.

**Figure 4:**
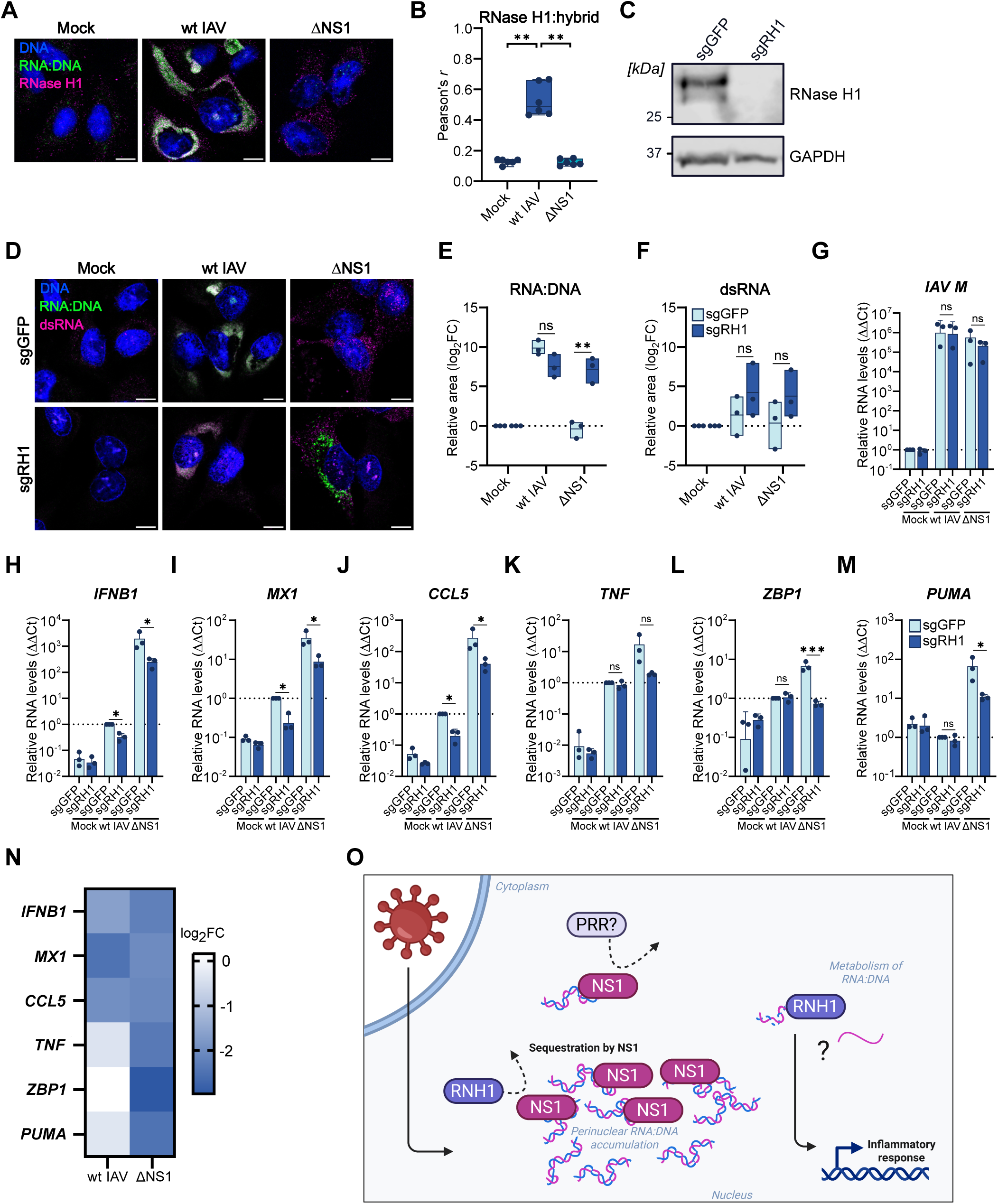
IAV NS1 protects RNA:DNA hybrids from the action of cellular RNase H1. **A-B**. Immunofluorescence microscopy analysis (A) and associated colocalization analysis (B) of A549 cells infected with wt IAV, IAV ΔNS1, or mock, at an MOI of 1 PFU/cell for 24 h, followed by methanol fixation, proteinase K treatment, and staining for RNA:DNA hybrids (S9.6; green), RNase H1 (magenta), and DNA (DAPI; blue). Colocalization between RNase H1 and RNA:DNA hybrids was assessed by determining the Pearson’s correlation coefficient (*r*) between green and magenta signals. Boxplots indicate the minimum to maximum range (whiskers) and mean (horizontal line) *r* values from n=3 independent experiments. Each dot represents the *r* value from a single field of view. Statistical significance was determined by Mann-Whitney U test (\*\**P□*≤□0.01). **C**. Immunoblot validation of generated RNase H1 knock-out (KO) A549 cells. Cell lysates from the generated RNase H1 knock-out (KO; sgRH1), or control (sgGFP), A549 cells were analyzed by immunoblotting for RNase H1 or GAPDH. Representative of n=3 independent experiments. **D**. Immunofluorescence microscopy analysis of RNase H1 knock-out (KO; sgRH1), or control (sgGFP), A549 cells infected with wt IAV, IAV ΔNS1, or mock, at an MOI of 1 PFU/cell for 24 h. Methanol fixed, proteinase K-treated cells were stained for RNA:DNA hybrids (S9.6; green), dsRNA (9D5; magenta), and DNA (DAPI; blue). Representative of n=3 independent experiments. **E-F**. Quantification of cytoplasmic RNA:DNA hybrid signal (E) and dsRNA signal (F) from the experiment described in D. Boxplots indicate the range and mean log_2_FC of the area fraction occupied by signal relative to mock-infected controls from n=3 independent experiments. Each dot represents the average log_2_FC from multiple fields imaged per well. Dotted line indicates no change relative to mock. Statistical significance between groups was determined by unpaired t-test with Welch’s correction for each infection condition \*\**P□*≤□0.01; ns non-significant). **G-M**. Gene expression analysis of the indicated genes in RNase H1 knock-out (KO; sgRH1), or control (sgGFP), A549 cells infected with wt IAV, IAV ΔNS1, or mock, at an MOI of 1 PFU/cell for 8 h followed by RNA extraction and RT-qPCR. (G) IAV M segment transcript levels were measured relative to mock-infected control cells. (H-M) Host genes were measured relative to wt IAV-infected control cells. Bars represent means, with standard deviation error bars from n=3 independent experiments, and statistical significance was determined by Welch’s unpaired t-test on ΔCt values (\**P□*≤□0.05; \*\*\**P□*≤□0.001; ns non-significant). **N**. Heatmap representing the magnitude of fold change (FC) reduction in gene expression for the targets shown in (H-M), calculated as the log_2_ ratio of KO to control transcript levels for each infected condition. A value of 0 indicates no change, while negative values reflect the degree of reduction. **O**. Hypothetical model. IAV infection leads to the accumulation of host and viral origin RNA:DNA hybrids, which are spatially sequestered by NS1 into a perinuclear localization. This sequestration limits access of canonical PRRs or RNase H1 to PAMP-like RNA:DNA hybrids. RNase H1 action appears to metabolize RNA:DNA hybrids and promote an innate inflammatory response. RNH1: RNase H1. For all microscopy images, scale bars represent 10 µm.

### RNase H1 contributes to IAV-triggered host responses

We reasoned that RNase H1 may possess an antiviral role relating to its metabolization of RNA:DNA hybrids during infection if indeed the IAV NS1 protein limits this process. To address this possibility, we therefore investigated host transcriptional immune responses in both sgRH1 and sgGFP cells infected with wt IAV or IAV ΔNS1 at an MOI of 1 PFU/cell for 8h. Under these conditions, viral infection levels were comparable between the two cell-lines as assessed by RT-qPCR for viral M RNA (**Fig. 4G**). Nevertheless, it was striking to observe reduced expression of the innate immunity related genes, *IFNB1* and *MX1*, as well as the inflammation-related genes, *CCL5*, *TNF*, *ZBP1* and *PUMA*, in the infected sgRH1 cell population as compared with the infected sgGFP cells (**Figs. 4H-M**). This suggests that RNase H1 promotes host innate responses during IAV infection, an unexpected possibility given that high levels of RNA:DNA hybrids have been associated with exacerbated innate immune activation (27, 28, 45). Interestingly, specifically for *TNF*, *ZBP1* and *PUMA* (genes primarily involved in inflammation, innate immune signaling and apoptosis), reduced expression in sgRH1 cells was most evident during infection with IAV ΔNS1, and not wt IAV (**Figs. 4K-N**). This distinction between groups of genes might be related to the regulatory mechanisms governing their expression: *IFNB1*, *MX1* and *CCL5* are part of the early antiviral response, primarily controlled by IFN-regulatory factor 3 (IRF3) and nuclear factor-ĸB (NF-ĸB) (46, 47); while *ZBP1*, *TNF* and *PUMA* are components of the PANoptosis pathway that is largely regulated by IRF1 and NF-ĸB (48), and which is a later response driven by the hyper-stimulated IFN-I-signaling typical of IAV ΔNS1 infections (49). Overall, our data are consistent with a hypothetical model whereby IAV infection-induced RNA:DNA hybrids of cellular and viral origins are metabolized by RNase H1 to promote stimulation of an inflammatory-dominated host response (**Fig. 4O**). This unanticipated function of RNase H1 may be via an unknown direct function of the protein, or perhaps via the generation of immunogenic cytosolic single-stranded DNA, the by-product of ribonuclease action. However, in the presence of IAV NS1, the viral protein appears to aid sequestration of RNA:DNA hybrids into perinuclear regions and to limit functional access by RNase H1, thus dampening RNase H1-mediated host responses.

## Discussion

While RNA:DNA hybrids are established modulators of cellular genome integrity and innate immunity, and serve as essential replication intermediates for retroviruses and contributors to the replication of certain DNA viruses, their presence and significance in RNA virus infections have remained largely unexplored. Here, our study provides evidence that IAV infection triggers an unexpected accumulation of perinuclear RNA:DNA hybrids of cellular and viral origin. This is notable as IAV is an RNA virus that replicates in the absence of a DNA intermediate. The mechanism underlying the generation of such RNA:DNA hybrids is therefore intriguing, but given that IAV infection is well-characterized to trigger the upregulation of TEs (13, 18, 50), which can encode endogenous reverse transcriptases (51), it seems plausible that cellular reverse transcriptases may at least contribute to non-specific IAV-sequence origin hybrid formation, akin to observations with other virus infections (22–25). In addition, cellular RNA:DNA hybrids can arise during RNA polymerase II-mediated stresses such as transcription-replication conflicts and RNA polymerase II stalling (26–28), which are described to be activated by IAV infections (16, 31). Thus, the appearance of RNA:DNA hybrids during IAV infection may be a general signal (or PAMP) to the cell, warning of an abnormal situation that has to be resolved.

A critical observation is that the IAV NS1 protein facilitates the presence and spatial sequestration of infection-induced RNA:DNA hybrids, effectively shielding them from metabolism by RNase H1. While NS1 is well-characterized as a dsRNA-binding immune-antagonist protein (41, 52), its canonical N-terminal RNA-binding domain does not interact with high affinity to RNA:DNA hybrids (53). Nevertheless, there is a report that NS1 can engage with dsDNA (54), and it is known that C-terminal domain interactions of NS1 leading to its multimerization can increase the ability of NS1 to engage with nucleic acid species (52, 55). Thus, we speculate that the direct or indirect interaction of NS1 with RNA:DNA hybrids may constitute a strategy to limit the action of RNase H1 activity. Importantly, our experiments suggest that RNase H1 normally contributes to IAV-triggered antiviral responses, including type I IFN and inflammatory gene expression programs. Therefore, while NS1 appears to counterintuitively stabilize potentially immunostimulatory RNA:DNA hybrids, which have previously been shown to bind and activate innate PRRs such as cGAS, TLR3, and ZBP1 (27, 30), its ability to limit metabolic access to these hybrids by RNase H1 seems key to preventing the activation of a previously unappreciated RNase H1-mediated host defense pathway. These actions of NS1 may not be mutually exclusive, and its sequestration of RNA:DNA hybrids into perinuclear regions may also shield them from directly activating canonical PRRs.

The precise molecular mechanism by which RNase H1-mediated metabolism of infection-induced RNA:DNA hybrids stimulates a host antiviral response remains intriguing. *A priori*, it may be that RNase H1 functions similarly to classical PRR sensors, thereby acting to detect abnormal RNA:DNA hybrids to trigger a downstream response in a yet unknown activity that may be enabled by its metabolic processing of the hybrids. While RNase H1 canonically functions to resolve RNA:DNA hybrid stress (43, 56), which would typically be thought to limit the accumulation of immunostimulatory hybrids, it is also possible that the metabolic activity of RNase H1 generates specific immunogenic products, including shorter more accessible hybrids or ssDNA that might be sensed and acted upon by classical PRRs. In this regard, persistent cellular DNA damage can result in RNA:DNA hybrid containing R-loop formation, which subsequently results in the cytoplasmic accumulation of ssDNA and the stimulation of pro-inflammatory responses (57). Thus, a similar mechanism may occur with IAV-induced RNA:DNA hybrids that escape sequestration by the viral NS1 protein, whereby RNase H1 acts as an intermediary to generate cytoplasmic pro-inflammatory ssDNA. In this context, the metabolic processing of RNA:DNA hybrids, rather than their presence alone, may dictate the magnitude of downstream responses. Identifying the specific signaling nodes that govern this putative antiviral pathway will be essential, and beyond understanding basic biology could reveal new insights into events underlying protection from infection or exacerbated deleterious pro-inflammatory signaling.

Taken together, our work uncovers an unexpected role for host and viral origin RNA:DNA hybrids in IAV infection biology, and suggests unanticipated cellular defense mechanisms that are targeted for viral antagonism. These observations contribute to understanding the complexity of host nucleic acid sensing in the self versus non-self paradox of immunity, and could provide the future basis for investigating new therapeutic interventions to strengthen antiviral resistance or to suppress aberrant autoinflammation.

## Methods

### Cells and viruses

A549, HEK293T, Vero CCL-81 and MDCK cells were cultured in Dulbecco’s Modified Eagle’s Medium (DMEM) (Life Technologies) supplemented with 10% (vol/vol) fetal calf serum (FCS), 100 units/mL penicillin, and 100 μg/mL streptomycin (Gibco Life Technologies). MRC5 cells were cultured in Minimum Essential Medium Eagle (MEM) (Sigma-Aldrich) supplemented with 10% (vol/vol) fetal bovine serum (FBS), 100 units/mL penicillin, 100□μg/mL streptomycin, 2□mM GlutaMAX (Thermo Fisher), and 1% non-essential amino acids (NEAA; Thermo Fisher). Calu-3 cells were cultured in DMEM supplemented with 20% (vol/vol) FBS, 100 units/mL penicillin, and 100 μg/mL streptomycin. IAV strain A/WSN/1933 (H1N1) was propagated in MDCK cells as described (58). The NS1-deficient WSN/33 variant (IAV ΔNS1), kindly provided by Balaji Manicassamy (University of Iowa, USA) (59), was propagated in MDCK cells stably expressing the NS1 protein from IAV strain A/PR8/34 (58) and the NPro protein from Bovine Viral Diarrhea Virus (60). The sequences of both IAV strains were verified by next-generation sequencing, and metagenomic analysis (https://github.com/medvir/virome-protocols) was performed to confirm the absence of any contaminating viral pathogens. A/Brisbane/59/2007 (H1N1), A/Brisbane/10/2007 (H3N2), A/Duck/Alberta/35/1976 (H1N1), A/Duck/Ukraine/1/1963 (H3N8), and A/Puerto Rico/8/34 (PR8; H1N1) working stocks were kindly provided by Silke Stertz (University of Zurich, Switzerland). rRSV-A2-GFP5 (R125, ViraTree; strain A2), PIV2-GFP (P222, ViraTree; strain V94), PIV5-GFP3 (P523, ViraTree; strain W3A) and MeV-GFP (OV1006, Imanis) were propagated in Vero CCL-81 cells and titrated as described previously (61). HSV-1-GFP (C-12) was kindly provided by Stacey Efstathiou (University of Cambridge, UK) (62), and was also propagated in Vero CCL-81 cells. A working stock of SFV was kindly provided by Jovan Pavlovic (University of Zurich, Switzerland).

### Virus infections

Cells were seeded at 4 x 10^6^ cells per 10 cm culture dish or 1 x 10^5^ cells per well of a 24-well plate and infected the next day at the indicated multiplicity of infection (MOI). For infections with PIV2, PIV5, MeV, RSV, and HSV-1, the inoculum was prepared in DMEM supplemented with 2% FBS, 100 units/mL penicillin, and 100 μg/mL streptomycin. Cells were then incubated with the inoculum for 1 h at 37°C, washed with PBS, and then overlaid with DMEM supplemented with 2% FBS, 100 units/mL penicillin, and 100 μg/mL streptomycin. For infections with IAV and SFV, inoculum was prepared in PBS containing 0.3% BSA, 1 mM Ca^2+^/Mg^2+^, 100 units/mL penicillin, and 100 μg/mL streptomycin. Cells were incubated with the inoculum for 1 h at 37 °C, washed with PBS, and then overlaid with DMEM supplemented with 0.1% FBS, 0.3% BSA, 20 mM HEPES, 100 units/mL penicillin, and 100 μg/mL streptomycin.

### Immunofluorescence confocal microscopy

Cells were seeded on glass coverslips in 24-well plates and infected with the indicated viruses as described above. At the time of collection, cells were washed twice with PBS, fixed and permeabilized with ice-cold methanol (MeOH) for 15 min at -20□°C, and washed four times with 1 mL of ice-cold PBS, followed by incubation with 0.01□U/mL Proteinase K in 50□mM Tris-HCl (pH 8.0) and 5□mM CaCl_2_ for 30□min at 37□°C. After washing three times with PBS, blocking was performed with 2% FBS in PBS (500□µL/well) for 30-60□min at room temperature (RT). Primary antibody staining was performed in PBS with 2% FBS using the following antibodies at the indicated dilutions: mouse monoclonal anti-RNA:DNA hybrid [S9.6] (ENH001, Kerafast, 1:1000), rabbit monoclonal anti-dsRNA [9D5] (Ab00458-23.0, Absolute Antibody; 1:1000), mouse monoclonal anti-dsRNA [9D5] (Ab00458-1.1, Absolute Antibody; 1:1000), rabbit polyclonal anti-RNASEH1 (15606-1-AP, Proteintech, 1:500), rabbit polyclonal anti-NS1 (PA5-32243, Invitrogen; 1:100), rabbit polyclonal anti-NP (kind gift from Jovan Pavlovic, University of Zurich, Switzerland; 1:1000), or rabbit polyclonal anti-PB2 (kind gift from Peter Palese, Icahn School of Medicine at Mount Sinai, USA, 1:1000). After overnight incubation at 4°C, cells were washed three times with PBS and incubated for 1 h at RT in the dark with the appropriate secondary antibody diluted in PBS with 2% FBS and DAPI (10236276001, Sigma-Aldrich; 1:1000). The secondary antibodies used were donkey anti-mouse IgG Alexa Fluor 488 (A21202, Thermo Fisher; 1:1000) and donkey anti-rabbit IgG Alexa Fluor 555 (A-21428, Thermo Fisher; 1:1000). Cells were then washed three times with PBS and two times with distilled water. Coverslips were mounted with ProLong Gold AntiFade (P36930, Thermo Fisher), and imaging was performed using a Leica SP8 confocal microscope (Leica) and LasX (Leica) software.

For the detection of GFP, SFV C protein and IAV NP, an alternative protocol was used. Briefly, cells were fixed with 4% PFA for 15 min at RT and permeabilized/blocked for 1 h with confocal buffer (PBS supplemented with 50□mM ammonium chloride, 0.1% saponin, and 2% BSA) at RT. Primary antibodies were then diluted in confocal buffer and incubated with the samples overnight at 4°C. The following primary antibodies were used: rabbit anti-GFP (GTX113617, GeneTex, 1:1000), rabbit anti-SFV-C (kind gift from Jovan Pavlovic, University of Zurich, Switzerland; 1:200), or rabbit polyclonal anti-NP (kind gift from Jovan Pavlovic, University of Zurich, Switzerland; 1:1000). Following overnight incubation at 4°C, cells were washed three times with PBS and incubated for 1 h at RT in the dark with donkey anti-rabbit IgG Alexa Fluor 555 (A-21428, Thermo Fisher; 1:1000) and DAPI (10236276001, Sigma-Aldrich; 1:1000). Image acquisition was as described above. Images were subsequently processed using ImageJ/Fiji software and Python.

### Image analysis and quantification

All image analyses were implemented in Python (v3.10.9) using the custom pipelines described in the **Supplementary Information**. For intracellular signal quantification, nuclei were identified via the DAPI channel to seed a watershed expansion, defining the cytosolic area within a 35-pixel radius. Within this region, signal was identified using a replicate-specific threshold, and the resulting area fraction was normalized as a fold change (FC) relative to the mock average. To ensure robustness, the limit of detection (LoD) was used as a denominator if the mock value fell below this threshold. The entire workflow is provided in the **Supplementary Information** as **Supplementary Code 1: Cytoplasmic Signal**. For pixel-based colocalization, images were first pre-processed using global min-max intensity normalization to standardize dynamic ranges. Spatial correlation between markers was then quantified by calculating the Pearson correlation coefficient (*r*) across pixel intensities as detailed in the **Supplementary Information** as **Supplementary Code 2: Colocalization**.

### Immunoprecipitations

A549 cells were washed once with PBS and lysed for 30 min on ice in co-immunopreciptation (co-IP) lysis buffer (50□mM Tris-HCl pH 7.5, 150 mM NaCl, 1□mM EDTA, 1% Triton X-100) freshly supplemented with cOmplete Protease Inhibitor Cocktail (11836170; Roche) and RNasin® Ribonuclease Inhibitor (N2615; Promega) at 1:1000 dilution. Samples were collected using a cell scraper and sonicated on ice followed by centrifugation at 16,000 x g for 15 min at 4°C. Supernatants were incubated overnight at 4°C with constant rotation in the presence of 2 µg of primary antibody. Antibodies used included mouse anti-V5 monoclonal antibody (MCA1360; BioRad), mouse monoclonal anti-RNA:DNA hybrid [S9.6] (ENH001, Kerafast, 1:1000) and mouse monoclonal anti-dsRNA [9D5] (Ab00458-1.1, Absolute Antibody; 1:1000). Protein G Dynabeads (10004D; Thermo Fisher) were added for 2 h at 4°C with constant rotation to capture immune complexes. The beads were then washed five times with co-IP lysis buffer and bound proteins eluted using 2x Laemmli protein sample buffer (Bio-Rad) containing 10% β-mercaptoethanol. Total cell lysates and immunoprecipitated fractions were analyzed by immunoblotting.

### Immunoblotting

Cells were lysed in urea disruption buffer (6M urea, 2M β-mercaptoethanol, 4% SDS, bromophenol blue) followed by sonication to shear nucleic acids. Samples were heated at 95°C for 5□min and separated via SDS-PAGE on Bolt 4-12% Bis-Tris Plus Mini Protein Gels (Invitrogen). Proteins were then transferred to 0.45 μm nitrocellulose membranes (Amersham) using the Invitrogen Mini Gel Tank and Blot Module. Membranes were blocked in 5% milk-TBS-T (5% milk in tris-buffered saline, TBS, supplemented with 0.1% Tween 20), and incubated with the following primary antibodies: rabbit anti-RNASEH1 polyclonal antibody (15606-1-AP, Proteintech); rabbit anti-NS1 polyclonal antibody (PA5-32243, Invitrogen), or mouse anti-GAPDH monoclonal antibody (0411)(#sc-47724, Santa Cruz). After washing with TBS-T, membranes were incubated in 5% milk in TBS-T or 5% BSA-TBS-T with the appropriate secondary antibodies: IRDye 800CW goat anti-mouse IgG (926–32210, LI-COR Biosciences), IRDye 800CW goat anti-rabbit IgG (926–32211, LI-COR Biosciences), IRDye 680RD goat anti-mouse IgG (926–68070, LI-COR Biosciences), IRDye 680RD goat anti-rabbit IgG (926–68071, LI-COR Biosciences), or HRP-conjugated goat anti-rabbit IgG (H+L) (SA00001-2, Proteintech). Membranes were imaged with the Odyssey Fc Imager (LI-COR Biosciences).

### RNA extraction and RT-qPCR

RNA was isolated with the ReliaPrep RNA Cell Miniprep kit (Promega) according to the manufacturer’s protocol. 1 μg of RNA was reverse-transcribed into cDNA using the SuperScript IV First-Strand Synthesis System (18091300, Thermo Fisher) with an Oligo(dT)_15_ primer (Promega). qPCR was performed with the PowerTrack SYBR Green Master Mix (A46109, Thermo Fisher) on an ABI7300 Real-Time PCR system (Applied Biosystems) in technical duplicates. Data were processed using SDS Shell, and relative cDNA quantities were calculated according to the ΔΔCt method in relation to 18S rRNA. qPCR primers used for specific targets are listed in **Supplementary Table 1.**

### Generation of *RNASEH1* knockout cells

To generate sgRH1 (*RNASEH1* knockout) and sgGFP control cells, three specific single-guide RNAs (sgRNAs) targeting the *RNASEH1* locus and a non-targeting sgRNA targeting GFP (**Supplementary Table 2**) were cloned into the lentiCRISPRv2 vector (Addgene plasmid #52961(63); gift from Feng Zhang, Massachusetts Institute of Technology, USA). Lentiviral particles for sgRH1 cells were produced in HEK293T cells by co-transfecting three individual lentiCRISPRv2 plasmids (each containing one of the three sgRNAs; 1.8 µg) together with pMD2.G (Addgene plasmid #12259) and psPAX2 (each 1 µg; Addgene plasmids #12260, gifts from Didier Trono, EPFL, Switzerland) with FuGENE HD transfection reagent (Promega). Lentiviral particles for sgGFP cells were similarly produced using the single lentiCRISPRv2 plasmid. Seventy two hours later, lentivirus-containing supernatants were collected and clarified by low-speed centrifugation and filtration through a 0.45 μm filter prior to storage at -80°C. A549 cells were then transduced with the lentivirus-containing supernatants in the presence of polybrene (8 µg/mL, Sigma-Aldrich) and were selected as a single polyclonal pool with puromycin (1 µg/mL, Thermo Fisher) for at least 14 days before functional analyses.

### Cytoplasmic DNA:RNA immunoprecipitation (cytoDRIP)

cytoDRIP was performed as described previously (27), with minor modifications to the fractionation (18). Briefly, 4x10^6^ A549 cells were seeded in 10 cm dishes and infected the next day for 24 h. Cells were then collected with trypsinization, centrifuged at 1200 RPM for 2 min, and the cell pellet washed once with PBS. Cells were lysed on ice for 5 min in 200 μL cell lysis buffer (10mM Tris pH 7.4, 150 mM NaCl, 0.15% IGEPAL CA-630), prepared and sterile filtered through a Steritop-GP 0.22 μm filter unit in advance, and freshly supplemented with cOmplete Protease Inhibitor Cocktail (11836170; Roche) and RNasin® Ribonuclease Inhibitor (N2615; Promega) at 1:1000 dilution immediately before use. Lysates were gently layered onto 500 μL of ice-cold sucrose buffer (10 mM Tris pH 7.4, 150 mM NaCl, 24% sucrose), also sterile-filtered in advance and freshly supplemented with RNasin® (1:1000), in protein LoBind 1.5 mL tubes before centrifugation at 3,500 x g for 10 min. The cytoplasmic supernatant was further cleared by centrifugation at 14,000 x g for 1 min in a fresh tube and collected. A small aliquot (5%) of this fraction was reserved for RNA isolation using the ReliaPrep RNA Cell Miniprep kit (Promega) to assess the purity of the cytoplasmic fraction. The nuclear fraction was briefly washed with 1 mL ice-cold PBS-EDTA (1x PBS, 500 μM EDTA pH 8.0; freshly supplemented with RNasin® at 1:1000), followed by centrifugation at 3500 x g for 1 min. The PBS-EDTA was then carefully removed from the nucleus-associated pellet. For RNA isolation from this fraction, the pellet was lysed in BL+TG buffer from the ReliaPrep RNA Cell Miniprep kit (Promega). The cytoplasmic fraction was then incubated in 0.4% SDS and 8 μL proteinase K (3115836001; Sigma-Aldrich) for 1 h at 37°C. DNA was extracted by phenol:chloroform:isoamyl alcohol (25:24:1) (A156.3; Roth), followed by ethanol precipitation. As indicated, samples were digested for 4-6 h at 37°C with 80 U RNase H (M0297; NEB) in 1x NEB RNase H buffer followed by DNA extraction and ethanol precipitation. For immunoprecipitation, 10 μL Protein G Dynabeads (10004D; Thermo Fisher) were incubated with 10 μg of mouse monoclonal anti-RNA:DNA hybrid [S9.6] antibody (ENH001; Kerafast) or mouse anti-V5 antibody (MCA1360; Bio-Rad) in 1x binding buffer (20□mM Tris-HCl pH□8.0, 2 mM EDTA, 1% Triton X-100, 150 mM NaCl, 0.5% sodium deoxycholate) for 4-6□h at 4°C. Simultaneously, the DNA extracts were precleared with 5 μL Protein G Dynabeads in 1x binding buffer for 1-2 h at 4°C. The precleared samples were then incubated overnight with the beads coated with S9.6 or anti-V5 antibodies at 4°C with constant rotation. Beads were subsequently washed with Tris/Saline/EDTA (TSE) buffer (20 mM Tris-HCl, pH 8.0; 2 mM EDTA; 1% Triton X-100; 0.1% SDS; 150 mM NaCl) followed by Tris-EDTA (TE) buffer. Bound complexes were then eluted in 250 μL elution buffer (50 mM Tris, pH 8; 10 mM EDTA; 0.5% SDS) and 8 μL proteinase K (20 mg/mL) for 50 mins at 50°C. DNA was extracted by phenol:chloroform:isoamyl alcohol (25:24:1), followed by ethanol precipitation.

### Library preparation and sequencing for cytoDRIP-seq

Second-strand synthesis was performed on ssDNA using random hexamers, dNTPs, and the Large (Klenow) Fragment (NEB) according to the manufacturer’s instructions, followed by purification with 2x volume AMPure XP beads (Beckman Coulter). Sequencing libraries were prepared from the resulting dsDNA using the Nextera XT DNA Library Preparation Kit and sequenced on the MiSeq platform (Illumina, to a depth of ≥1^5^ single-end 151-bp reads).

### cytoDRIP-seq data analysis

Raw sequencing reads were initially processed using fastp v0.23.4 for quality control and adapter trimming (64). Reads were mapped to a concatenation of the human (GRCh38.p14) and influenza A virus (ASM3791505v1) genomes using Bowtie2 v2.5.4 with default parameters (65). Reads that aligned to TE consensus sequences using Bowtie2 in local mode to allow for soft clipping were separately identified. Duplicated reads were removed using Picard MarkDuplicates v3.2.0 (https://github.com/broadinstitute/picard). Only human chromosomes were considered for further downstream analyses; unplaced contigs/scaffolds were discarded. As the read counts for viral segments were orders of magnitude lower than those aligning to human chromosomes, this category was treated separately. Using the Hmisc R package, we divided the read counts for viral segments into three quantiles (High/Medium/Low levels) and retained only the counts categorized as High (i.e., we discarded as noise the signal based on less than ∼100 reads).

The bamCoverage tool from the deepTools suite v.3.5.6 was used to generate sequencing depth-normalized coverage signal tracks (CPM) with bin size = 1 bp (66). Subtracted signals (RNase H treatment subtracted from mock-treated samples) were generated using the bamCompare tool from the deepTools suite with CPM normalization, bin size = 1 bp, and smooth length = 5. For visualization purposes, instances of negative signal were capped to zero. To quantify the sequencing depth-normalized signal spanning genomic regions of different length (e.g., whole chromosomes, viral segments, or genomic features) we computed mean signal density per base pair using bedtools intersect v2.31.1 and multiplied the base-pair overlap of the signal track by its corresponding intensity (67). The cumulative signal was then divided by total length of the feature of interest to obtain length-normalized mean signal density (average CPM per base pair). Human intronic and intergenic genomic features were computed using GTFtools v0.9.0 (68). Pseudocounts, log scaling, or ratio between signals were applied when appropriate for visualization purposes and downstream comparative analyses. Unless specified otherwise, base R (v4.5.3) was used for statistical analyses, and the tidyverse and ggplot2 (https://ggplot2.tidyverse.org) R packages were used for data transformation and visualization, respectively (69).

Further bioinformatic analysis was performed using VirMet (https://github.com/medvir/VirMet) for viral reads, in which reads were filtered to exclude low-quality sequences (Phred score < 20), short reads (<75 bp), and low-entropy reads (complexity < 70%, as previously defined (70)). Quality-filtered reads were aligned to the influenza A virus genome (A/WSN/1933 TS61 (H1N1)) using Burrows-Wheeler Aligner (v0.7.17, (71)) and processed with SAMtools (v1.10, (72)). Sequencing depth was normalized to reads per million (RPM) to enable comparisons across samples.

### Statistical analysis

Statistical analyses were primarily performed using GraphPad Prism 7. The specific statistical tests applied, along with their corresponding *P* values are indicated in the figure legends. RT-qPCR data were analyzed using the 2^−ΔΔCT^ method as described previously (73).

### Graphics

Objects in figures 2A, and 4O were created in BioRender. Kwaschik, F (2026): https://BioRender.com/hyelh6g; https://BioRender.com/wjluz15.

### Data availability

The unique datasets produced in this study will be deposited to the Gene Expression Omnibus database (GEO) following peer review.

## Supporting information

Supplementary Figure Legends and Codes

## Acknowledgements

We thank Balaji Manicassamy, Silke Stertz, Stacey Efstathiou, Peter Palese, and Jovan Pavlovic for viruses and antibodies. We also thank Feng Zhang and Didier Trono for plasmids via Addgene. We are grateful to Massimo Lopes, Renate König and Liam Childs for preliminary technical consultations, and to Magdalena Crossley for providing the cytoDRIP-seq protocol and guidance. We also thank Stefan Schmutz for initial technical assistance. The research leading to these results received funding from the Swiss National Science Foundation (www.snf.ch; grant 310030_214957 to BGH) and the University of Zurich (www.uzh.ch; Forschungskredit grant FK-25-030 to FK). The funders had no role in study design, data collection, data interpretation, or the decision to submit the work for publication.

## References

1. Bartok E, Hartmann G. Immune Sensing Mechanisms that Discriminate Self from Altered Self and Foreign Nucleic Acids. Immunity. 2020;53(1):54–77.

2. Chiang JJ, Sparrer KMJ, van Gent M, Lassig C, Huang T, Osterrieder N, et al. Viral unmasking of cellular 5S rRNA pseudogene transcripts induces RIG-I-mediated immunity. Nat Immunol. 2018;19(1):53–62.

3. Moriyama M, Koshiba T, Ichinohe T. Influenza A virus M2 protein triggers mitochondrial DNA-mediated antiviral immune responses. Nat Commun. 2019;10(1):4624.

4. Ariav Y, Ch’ng JH, Christofk HR, Ron-Harel N, Erez A. Targeting nucleotide metabolism as the nexus of viral infections, cancer, and the immune response. Sci Adv. 2021;7(21):eabg6165.

5. Chen Z, Behrendt R, Wild L, Schlee M, Bode C. Cytosolic nucleic acid sensing as driver of critical illness: mechanisms and advances in therapy. Signal Transduction and Targeted Therapy. 2025;10(1):90.

6. Hu MM, Shu HB. Cytoplasmic Mechanisms of Recognition and Defense of Microbial Nucleic Acids. Annu Rev Cell Dev Biol. 2018;34:357–79.

7. Chiappinelli KB, Strissel PL, Desrichard A, Li H, Henke C, Akman B, et al. Inhibiting DNA Methylation Causes an Interferon Response in Cancer via dsRNA Including Endogenous Retroviruses. Cell. 2015;162(5):974–86.

8. Cuellar TL, Herzner AM, Zhang X, Goyal Y, Watanabe C, Friedman BA, et al. Silencing of retrotransposons by SETDB1 inhibits the interferon response in acute myeloid leukemia. J Cell Biol. 2017;216(11):3535–49.

9. Roulois D, Loo Yau H, Singhania R, Wang Y, Danesh A, Shen SY, et al. DNA-Demethylating Agents Target Colorectal Cancer Cells by Inducing Viral Mimicry by Endogenous Transcripts. Cell. 2015;162(5):961–73.

10. Thomas CA, Tejwani L, Trujillo CA, Negraes PD, Herai RH, Mesci P, et al. Modeling of TREX1-Dependent Autoimmune Disease using Human Stem Cells Highlights L1 Accumulation as a Source of Neuroinflammation. Cell Stem Cell. 2017;21(3):319-+.

11. Okude H, Ori D, Kawai T. Signaling Through Nucleic Acid Sensors and Their Roles in Inflammatory Diseases. Front Immunol. 2021;11:625833-.

12. Uggenti C, Lepelley A, Crow YJ. Self-Awareness: Nucleic Acid-Driven Inflammation and the Type I Interferonopathies. Annu Rev Immunol. 2019;37(Volume 37, 2019):247–67.

13. Schmidt N, Domingues P, Golebiowski F, Patzina C, Tatham MH, Hay RT, et al. An influenza virus-triggered SUMO switch orchestrates co-opted endogenous retroviruses to stimulate host antiviral immunity. Proc Natl Acad Sci U S A. 2019;116(35):17399–408.

14. Hale BG. Antiviral immunity triggered by infection-induced host transposable elements. Curr Opin Virol. 2022;52:211–6.

15. Marston JL, Greenig M, Singh M, Bendall ML, Duarte RRR, Feschotte C, et al. SARS-CoV-2 infection mediates differential expression of human endogenous retroviruses and long interspersed nuclear elements. JCI Insight. 2021;6(24).

16. Yin C, Fedorov A, Guo H, Crawford JC, Rousseau C, Zhong X, et al. Host cell Z-RNAs activate ZBP1 during virus infections. Nature. 2025;648(8094):707–16.

17. Cai C, Tang Y-D, Xu G, Zheng C. The crosstalk between viral RNA- and DNA-sensing mechanisms. Cellular and Molecular Life Sciences. 2021;78(23):7427–34.

18. Lork M, Childs L, Lieber G, Kwaschik F, Konig R, Hale BG. Regulated localization of transposable element RNA during influenza A virus infection. EMBO Rep. 2025;26(14):3506–28.

19. Nakano M, Miyamoto S, Ohnishi C, Nogami C, Hirose N, Fujita-Fujiharu Y, et al. Influenza A virus circumvents the innate immune response through the sequestration of double-stranded RNA. J Virol. 2025;99(10):e0073725.

20. Oade MS, te Velthuis AJW. Molecular Insights into Noncanonical Influenza Virus Replication and Transcription. Annual Review of Virology. 2025;12(Volume 12, 2025):259–76.

21. Sadeq S, Al-Hashimi S, Cusack CM, Werner A. Endogenous Double-Stranded RNA. Non-Coding RNA [Internet]. 2021; 7(1):[15 p.].

22. Wu J, Wu C, Xing F, Cao L, Zeng W, Guo L, et al. Endogenous reverse transcriptase and RNase H-mediated antiviral mechanism in embryonic stem cells. Cell Res. 2021;31(9):998–1010.

23. Klenerman P, Hengartner H, Zinkernagel RM. A non-retroviral RNA virus persists in DNA form. Nature. 1997;390(6657):298–301.

24. Geuking MB, Weber J, Dewannieux M, Gorelik E, Heidmann T, Hengartner H, et al. Recombination of retrotransposon and exogenous RNA virus results in nonretroviral cDNA integration. Science. 2009;323(5912):393–6.

25. Zhang LG, Richards A, Barrasa MI, Hughes SH, Young RA, Jaenisch R. Reverse-transcribed SARS-CoV-2 RNA can integrate into the genome of cultured human cells and can be expressed in patient-derived tissues. P Natl Acad Sci USA. 2021;118(21):e2105968118-e.

26. Niehrs C, Luke B. Regulatory R-loops as facilitators of gene expression and genome stability. Nat Rev Mol Cell Biol. 2020;21(3):167–78.

27. Crossley MP, Song C, Bocek MJ, Choi JH, Kousouros JN, Sathirachinda A, et al. R-loop-derived cytoplasmic RNA-DNA hybrids activate an immune response. Nature. 2023;613(7942):187–94.

28. Crossley MP, Bocek M, Cimprich KA. R-Loops as Cellular Regulators and Genomic Threats. Mol Cell. 2019;73(3):398–411.

29. Rigby RE, Webb LM, Mackenzie KJ, Li Y, Leitch A, Reijns MA, et al. RNA:DNA hybrids are a novel molecular pattern sensed by TLR9. EMBO J. 2014;33(6):542–58.

30. He J, Zhu Y, Tian Z, Liu M, Gao A, Fu W, et al. ZBP1 senses spliceosome stress through Z-RNA:DNA hybrid recognition. Mol Cell. 2025;85(9):1790–805 e7.

31. Bauer DLV, Tellier M, Martinez-Alonso M, Nojima T, Proudfoot NJ, Murphy S, et al. Influenza Virus Mounts a Two-Pronged Attack on Host RNA Polymerase II Transcription. Cell Rep. 2018;23(7):2119–29 e3.

32. Son KN, Liang ZG, Lipton HL. Double-Stranded RNA Is Detected by Immunofluorescence Analysis in RNA and DNA Virus Infections, Including Those by Negative-Stranded RNA Viruses. Journal of Virology. 2015;89(18):9383–92.

33. Boguslawski SJ, Smith DE, Michalak MA, Mickelson KE, Yehle CO, Patterson WL, et al. Characterization of monoclonal antibody to DNA.RNA and its application to immunodetection of hybrids. J Immunol Methods. 1986;89(1):123–30.

34. Smolka JA, Sanz LA, Hartono SR, Chedin F. Recognition of RNA by the S9.6 antibody creates pervasive artifacts when imaging RNA:DNA hybrids. J Cell Biol. 2021;220(6).

35. Pichlmair A, Schulz O, Tan CP, Rehwinkel J, Kato H, Takeuchi O, et al. Activation of MDA5 requires higher-order RNA structures generated during virus infection. J Virol. 2009;83(20):10761–9.

36. Weber F, Wagner V, Rasmussen SB, Hartmann R, Paludan SR. Double-Stranded RNA Is Produced by Positive-Strand RNA Viruses and DNA Viruses but Not in Detectable Amounts by Negative-Strand RNA Viruses. Journal of Virology. 2006;80(10):5059–64.

37. Eisfeld AJ, Neumann G, Kawaoka Y. At the centre: influenza A virus ribonucleoproteins. Nat Rev Microbiol. 2015;13(1):28–41.

38. Hutchinson JN, Ensminger AW, Clemson CM, Lynch CR, Lawrence JB, Chess A. A screen for nuclear transcripts identifies two linked noncoding RNAs associated with SC35 splicing domains. BMC Genomics. 2007;8(1):39.

39. Samacoits A, Chouaib R, Safieddine A, Traboulsi AM, Ouyang W, Zimmer C, et al. A computational framework to study sub-cellular RNA localization. Nat Commun. 2018;9(1):4584.

40. Sanz LA, Chedin F. High-resolution, strand-specific R-loop mapping via S9.6-based DNA-RNA immunoprecipitation and high-throughput sequencing. Nat Protoc. 2019;14(6):1734–55.

41. Hale BG, Randall RE, Ortin J, Jackson D. The multifunctional NS1 protein of influenza A viruses. J Gen Virol. 2008;89(Pt 10):2359–76.

42. Wahba L, Amon JD, Koshland D, Vuica-Ross M. RNase H and multiple RNA biogenesis factors cooperate to prevent RNA:DNA hybrids from generating genome instability. Mol Cell. 2011;44(6):978–88.

43. Lockhart A, Pires VB, Bento F, Kellner V, Luke-Glaser S, Yakoub G, et al. RNase H1 and H2 Are Differentially Regulated to Process RNA-DNA Hybrids. Cell Rep. 2019;29(9):2890–900 e5.

44. Sollier J, Cimprich KA. Breaking bad: R-loops and genome integrity. Trends Cell Biol. 2015;25(9):514–22.

45. Cristini A, Tellier M, Constantinescu F, Accalai C, Albulescu LO, Heiringhoff R, et al. RNase H2, mutated in Aicardi-Goutieres syndrome, resolves co-transcriptional R-loops to prevent DNA breaks and inflammation. Nat Commun. 2022;13(1):2961.

46. McNab F, Mayer-Barber K, Sher A, Wack A, O’Garra A. Type I interferons in infectious disease. Nat Rev Immunol. 2015;15(2):87–103.

47. Hoffmann A, Levchenko A, Scott ML, Baltimore D. The IkappaB-NF-kappaB signaling module: temporal control and selective gene activation. Science. 2002;298(5596):1241–5.

48. Nguyen LN, Kanneganti TD. PANoptosis in Viral Infection: The Missing Puzzle Piece in the Cell Death Field. J Mol Biol. 2022;434(4):167249.

49. Forero A, Ozarkar S, Li H, Lee CH, Hemann EA, Nadjsombati MS, et al. Differential Activation of the Transcription Factor IRF1 Underlies the Distinct Immune Responses Elicited by Type I and Type III Interferons. Immunity. 2019;51(3):451–64 e6.

50. Chen X, Pacis A, Aracena KA, Gona S, Kwan T, Groza C, et al. Transposable elements are associated with the variable response to influenza infection. Cell Genom. 2023;3(5):100292.

51. Fukuda S, Varshney A, Fowler BJ, Wang SB, Narendran S, Ambati K, et al. Cytoplasmic synthesis of endogenous Alu complementary DNA via reverse transcription and implications in age-related macular degeneration. Proc Natl Acad Sci U S A. 2021;118(6):e2022751118-e.

52. Hale BG. Conformational plasticity of the influenza A virus NS1 protein. J Gen Virol. 2014;95(Pt 10):2099–105.

53. Chien CY, Xu Y, Xiao R, Aramini JM, Sahasrabudhe PV, Krug RM, et al. Biophysical characterization of the complex between double-stranded RNA and the N-terminal domain of the NS1 protein from influenza A virus: evidence for a novel RNA-binding mode. Biochemistry. 2004;43(7):1950–62.

54. Anastasina M, Le May N, Bugai A, Fu Y, Soderholm S, Gaelings L, et al. Influenza virus NS1 protein binds cellular DNA to block transcription of antiviral genes. Biochim Biophys Acta. 2016;1859(11):1440–8.

55. Kerry PS, Ayllon J, Taylor MA, Hass C, Lewis A, Garcia-Sastre A, et al. A transient homotypic interaction model for the influenza A virus NS1 protein effector domain. PLoS One. 2011;6(3):e17946.

56. Shen W, Sun H, De Hoyos CL, Bailey JK, Liang XH, Crooke ST. Dynamic nucleoplasmic and nucleolar localization of mammalian RNase H1 in response to RNAP I transcriptional R-loops. Nucleic Acids Res. 2017;45(18):10672–92.

57. Chatzidoukaki O, Stratigi K, Goulielmaki E, Niotis G, Akalestou-Clocher A, Gkirtzimanaki K, et al. R-loops trigger the release of cytoplasmic ssDNAs leading to chronic inflammation upon DNA damage. Sci Adv. 2021;7(47):eabj5769.

58. Hale BG, Jackson D, Chen YH, Lamb RA, Randall RE. Influenza A virus NS1 protein binds p85beta and activates phosphatidylinositol-3-kinase signaling. Proc Natl Acad Sci U S A. 2006;103(38):14194–9.

59. Manicassamy B, Manicassamy S, Belicha-Villanueva A, Pisanelli G, Pulendran B, Garcia-Sastre A. Analysis of in vivo dynamics of influenza virus infection in mice using a GFP reporter virus. Proc Natl Acad Sci U S A. 2010;107(25):11531–6.

60. Hilton L, Moganeradj K, Zhang G, Chen YH, Randall RE, McCauley JW, et al. The NPro product of bovine viral diarrhea virus inhibits DNA binding by interferon regulatory factor 3 and targets it for proteasomal degradation. J Virol. 2006;80(23):11723–32.

61. Lieber G, Kwaschik F, Lork M, Schmidt N, Hale BG. TRIM28 is a target for paramyxovirus V proteins. PLoS Pathog. 2025;21(9):e1013487.

62. Glauser DL, Seyffert M, Strasser R, Franchini M, Laimbacher AS, Dresch C, et al. Inhibition of herpes simplex virus type 1 replication by adeno-associated virus rep proteins depends on their combined DNA-binding and ATPase/helicase activities. J Virol. 2010;84(8):3808–24.

63. Sanjana NE, Shalem O, Zhang F. Improved vectors and genome-wide libraries for CRISPR screening. Nature Methods. 2014;11(8):783–4.

64. Chen S. Ultrafast one-pass FASTQ data preprocessing, quality control, and deduplication using fastp. Imeta. 2023;2(2):e107.

65. Langmead B, Salzberg SL. Fast gapped-read alignment with Bowtie 2. Nature Methods. 2012;9(4):357–9.

66. Ramírez F, Ryan DP, Grüning B, Bhardwaj V, Kilpert F, Richter AS, et al. deepTools2: a next generation web server for deep-sequencing data analysis. Nucleic Acids Research. 2016;44(W1):W160–W5.

67. Quinlan AR, Hall IM. BEDTools: a flexible suite of utilities for comparing genomic features. Bioinformatics. 2010;26(6):841–2.

68. Li H-D, Lin C-X, Zheng J. GTFtools: a software package for analyzing various features of gene models. Bioinformatics. 2022;38(20):4806–8.

69. Wickham H, Averick M, Bryan J, Chang W, McGowan LDA, François R, et al. Welcome to the Tidyverse. Journal of open source software. 2019;4(43):1686.

70. Schmieder R, Edwards R. Quality control and preprocessing of metagenomic datasets. Bioinformatics. 2011;27(6):863–4.

71. Li H. Aligning sequence reads, clone sequences and assembly contigs with BWA-MEM. arXiv: Genomics. 2013.

72. Danecek P, Bonfield JK, Liddle J, Marshall J, Ohan V, Pollard MO, et al. Twelve years of SAMtools and BCFtools. GigaScience. 2021;10(2):giab008.

73. Livak KJ, Schmittgen TD. Analysis of Relative Gene Expression Data Using Real-Time Quantitative PCR and the 2−ΔΔCT Method. Methods. 2001;25(4):402–8.

