## Supplementary Table 1 for "Influenza A virus NS1 sequesters RNA:DNA hybrids to evade RNase H1-dependent innate immunity"

Supplementary Table 1: qPCR primers

| Name | Orientation | Sequence (5’→3’) |
| --- | --- | --- |
| 18S rRNA | Forward | GGCCCTGTAATTGGAATGACTC |
| 18S rRNA | Reverse | CCAAGATCCAACTACGAGCTT |
| IAV M (WHO M30F2) | Forward | ATGAGYCTTYTAACCGAGGTCGAAACG |
| IAV M (WHO M264R3) | Reverse | TGGACAAANCGTCTACGCTGCAG |
| IFNB1 | Forward | CATTACCTGAAGGCCAAGGA |
| IFNB1 | Reverse | CAGCATCTGCTGGTTGAAGA |
| MX1 | Forward | AGACAAGGTTGTGGACGTGG |
| MX1 | Reverse | TTCCTCCAGCAGATCCCTGA |
| CCL5 | Forward | GCTGTCATCCTCATTGCTACTG |
| CCL5 | Reverse | TGGTGTAGAAATACTCCTTGATGTG |
| TNF | Forward | CCTCTCTCTAATCAGCCCTCTG |
| TNF | Reverse | GAGGACCTGGGAGTAGATGAG |
| ZBP1 | Forward | CAGCCTCGACCTAGCCCA |
| ZBP1 | Reverse | TGCAGGATTCTTTGTTCAAGGTG |
| PUMA | Forward | GGACTCAGCATCGGAAGGT |
| PUMA | Reverse | GCACCAGCACAACAGCCTTT |
| GAPDH | Forward | CTGGCGTCTTCACCACCATGG |
| GAPDH | Reverse | CATCACGCCACAGTTTCCCGG |
| MALAT1 | Forward | GAAGGAAGGAGCGCTAACGA |
| MALAT1 | Reverse | TACCAACCACTCGCTTTCCC |
