## Supplementary Table 2 for "Influenza A virus NS1 sequesters RNA:DNA hybrids to evade RNase H1-dependent innate immunity"

Supplementary Table 2: LentiCRISPR oligonucleotides

| Name | Forward primer (5’→3’) | Reverse primer (5’→3’) |
| --- | --- | --- |
| sgRH1_1 | CACCGAAAGCACATGAAGCCGAGCG | AAACCGCTCGGCTTCATGTGCTTTC |
| sgRH1_2 | CACCGCACTGGAGAGGCTTACCCAG | AAACCTGGGTAAGCCTCTCCAGTGC |
| sgRH1_3 | CACCGTTAACTTACAAAGGATGGCC | AAACGGCCATCCTTTGTAAGTTAAC |
| sgGFP | CACCGGGGCGAGGAGCTGTTCACCG | AAACCGGTGAACAGCTCCTCGCCCC |
