## Supplementary Figure Legends and Codes for "Influenza A virus NS1 sequesters RNA:DNA hybrids to evade RNase H1-dependent innate immunity"

**Supplementary Information File** (Kwaschik *et al*.)

Contains Supplementary Figure Legends (Figures S1-S4) and Supplementary Codes 1-2

**Supplementary Figure Legends**

**Figure S1: Induction of RNA:DNA hybrids and dsRNA across different human cell lines by various IAV strains**

**A**. Colocalization analysis of RNA:DNA hybrids with dsRNA in A549 cells infected with IAV, or mock, at an MOI of 1 PFU/cell for 24 h. Pearson’s correlation coefficient (*r*) was quantified from immunofluorescence confocal images staining for RNA:DNA hybrids (S9.6) and dsRNA (9D5). Boxplots indicate the minimum to maximum range (whiskers) and mean (horizontal line) *r* values from n=3 independent experiments. Each dot represents the *r* value from a single field of view. Statistical significance was determined by Mann-Whitney U test (*** *P* ≤ 0.001). Related to Figure 1A.

**B-D**. Immunofluorescence microscopy images of MRC5 (B), Calu-3 (C), and HT-29 (D) cells infected, or mock, with IAV at an MOI of 1 PFU/cell for 24 h. Cells were fixed and permeabilized with methanol, proteinase K treated, and then stained for RNA:DNA hybrids (S9.6; green), dsRNA (9D5; magenta), and DNA (DAPI; blue).

**E**. Immunofluorescence microscopy images of A549 cells infected, or mock, with IAVs A/WSN/33 (MOI of 1 PFU/cell) or A/Brisbane/59/2007 (MOI of 2.5 PFU/cell) for 24 h, or A/Brisbane/10/2007 (MOI of 10 PFU/cell), A/Duck/Alberta/35/1976 (MOI of 20 PFU/cell), and A/Duck/Ukraine/1/1963 (MOI of 20 PFU/cell) for 48 h. Upper row: cells were fixed and permeabilized with methanol, proteinase K treated, and then stained for RNA:DNA hybrids (S9.6; green), dsRNA (9D5; magenta), or DNA (DAPI; blue). Lower row: same treatments stained for IAV NP (red) and DNA (DAPI; blue).

**F**. Immunofluorescence microscopy images of A549 cells infected, or mock, with IAV A/Puerto Rico/8/1934 (A/PR8/34) at an MOI of 1 PFU/cell for 24 h. Cells were fixed and permeabilized with methanol, proteinase K treated, and then stained for RNA:DNA hybrids (S9.6; green), dsRNA (9D5; magenta), or DNA (DAPI; blue).

**G.**  Quantification of cytoplasmic RNA:DNA hybrids from experiments in E-F. Each data point represents the average log_2_ fold change (FC) of the area fraction occupied by signal relative to mock-infected controls from n=2 independent experiments. For replicates where mock or infected signal was not detected, FC was calculated relative to the limit of detection (LoD). WSN, A/WSN/33; B10, A/Brisbane/10/2007; B59, A/Brisbane/59/2007; DA, A/Duck/Alberta/35/1976; DU, A/Duck/Ukraine/1/1963; PR8, A/PR8/34.

For all panels, data are representative of at least n=2 independent experiments. For microscopy images, scale bars represent 10 µm.

**Figure S2: Extended ΔDRIP-seq analysis**

**A**. Validation of subcellular fractionation and IAV infection levels by RT-qPCR. Fractionated samples from the experiments described in Figure 2 were assessed for IAV M, GAPDH, and MALAT1 RNA levels, which acted as markers for viral load, cytosolic fraction, and nuclear fraction, respectively. Bars represent mean values and standard deviations from n=3 independent replicates (each dot represents one replicate). T: Total; N: Nucleus; C: Cytosol. Related to Figure 2.

**B-C**. Abundance of host- and viral-derived sequences. Percentage of total sequencing reads mapping to the human (B) and IAV (C) genomes. Data represent the composition of non-RNase H1-treated samples to illustrate the initial distribution of captured sequences. Bars represent the mean of n=3 independent biological replicates, with individual dots indicating the values for each replicate. Error bars represent standard deviations.

**D**. Bar plot showing background-subtracted (ΔDRIP-seq) signal density of reads mapping to the human genome across samples immunoprecipitated with anti-V5 or S9.6 antibodies. Data represent the mean signal density (normalized for sequencing depth and feature length) relative to the mock sample immunoprecipitated with anti-V5 (for visualization purposes), with error bars representing standard deviations. Individual dots represent values from n=3 biological replicates.

**E**. Bar plot showing background-subtracted (ΔDRIP-seq) signal density of reads mapping to the IAV genome across samples immunoprecipitated with the indicated antibodies. Data represent the mean signal density (normalized to sequencing depth and feature length). Individual dots represent values from n=3 biological replicates, with error bars representing standard deviations.

**F**. Bar plot showing the average enrichment of viral signal across the eight IAV segments. Data represent the mean signal density (ΔRPM normalized to segment length) of n=3 biological replicates. Bars represent the mean, individual dots indicate the values of each replicate, and error bars represent the standard deviation.

**Figure S3: Subcellular distribution of RNA:DNA hybrids and dsRNA with IAV NS1 or PB2**

**A**. Immunofluorescence microscopy analysis of A549 cells infected, or mock, with IAV at an MOI of 1 PFU/cell for 24 h, followed by methanol fixation, proteinase K treatment, and staining for RNA:DNA hybrids (S9.6; green), NS1 (magenta), and DNA (DAPI; blue). Merged images are presented in Figure 3A.

**B**. Detection of dsRNA (9D5; green), NS1 (magenta), and DNA (DAPI; blue) in A549 cells treated as detailed in A. Merged images are presented in Figure 3C.

**C**. Detection of RNA:DNA hybrids (S9.6; green), PB2 (magenta), and DNA (DAPI; blue) in A549 cells treated as detailed in A. Merged images are presented in Figure 3E.

**D**. Detection of dsRNA (9D5; green), PB2 (magenta), and DNA (DAPI; blue) in A549 cells treated as detailed in A. Merged images are presented in Figure 3G.

For all panels, data are representative of at least n=3 independent experiments. Scale bars represent 10 µm.

**Figure S4: Co-localization of RNA:DNA hybrids and dsRNA during IAV infection in wild-type and RNase H1-deficient cells**

**A-B**. Quantification of RNA:DNA hybrid and dsRNA colocalization in mock, wt IAV, and IAVΔNS1 infected control (sgGFP) A549 cells (A) or RNase H1 knock-out (KO; sgRH1) A549 cells (B). Infections and immunofluorescence microscopy were carried out as described in Figure 4D. Co-localization of RNA:DNA hybrids and dsRNA was assessed by determining the Pearson’s correlation coefficient (*r*) between green (RNA:DNA) and magenta (dsRNA) signals. Boxplots indicate the minimum to maximum range (whiskers) and mean (horizontal line) *r* values from n=3 independent experiments. Each dot represents the *r* value from a single field of view. Statistical significance was determined by Mann-Whitney U test (*****P* ≤ 0.0001).

**Supplementary Code 1: Cytoplasmic Signal**

Description: The following script was used to calculate the cytoplasmic area fraction

```
import pandas as pd
from pathlib import Path
import numpy as np
from skimage import io, filters, measure, morphology, segmentation
from scipy import ndimage as ndi
from datetime import datetime
import matplotlib
matplotlib.use('Agg')
import matplotlib.pyplot as plt

"""
Script for Analysis of Cytoplasmic Signal (RNA:DNA Hybrids and dsRNA)
Calculates total cytoplasmic area and area fraction above limit of detection (LoD)
"""

import pandas as pd
from pathlib import Path
import numpy as np
from skimage import io, filters, measure, morphology, segmentation
from scipy import ndimage as ndi
from datetime import datetime
import matplotlib
matplotlib.use('Agg')
import matplotlib.pyplot as plt
import gc

def run_global_load_analysis(folder_path, max_dist=35, sigma_ch2=7.0, sigma_ch3=5.0, min_cluster_size=2, cal_percentile=98.0):
    # --- 1. SETUP ---
    base_path = Path(folder_path)
    timestamp = datetime.now().strftime("%Y%m%d_%H%M%S")
    run_dir = base_path / f"Analysis_Run_{timestamp}_GlobalLoad"
    run_dir.mkdir(parents=True, exist_ok=True)

    preview_dir, hist_dir = run_dir / "QC_Previews", run_dir / "QC_Histograms"
    for d in [preview_dir, hist_dir]: d.mkdir(exist_ok=True)

    # --- 2. CALIBRATE & LoD ---
    TRUST_LIMIT_CH2, FALLBACK_CH2 = 30.0, 54.0          #may need to be adapted based on signal to noise ratio
    TRUST_LIMIT_CH3, FALLBACK_CH3 = 2.0, 15.0           #may need to be adapted based on signal to noise ratio

    print(f"--- STARTING ANALYSIS: {timestamp} ---")
    mock_files = [f for f in base_path.glob("**/[Mm]ock*/*ch00.tif") if "2nd" not in f.name.lower()]
    if not mock_files: mock_files = list(base_path.glob("**/*ch00.tif"))[:5]

    ch2_cals, ch3_cals = [], []
    for dapi_path in mock_files:
        img_ch2_cal = io.imread(str(dapi_path).replace("ch00.tif", "ch01.tif"), as_gray=True)
        ch2_cals.append(np.percentile(img_ch2_cal, cal_percentile))
        img_ch3_cal = io.imread(str(dapi_path).replace("ch00.tif", "ch02.tif"), as_gray=True)
        ch3_cals.append(np.percentile(img_ch3_cal, cal_percentile))

    auto_t2 = np.median(ch2_cals) * sigma_ch2
    auto_t3 = np.median(ch3_cals) * sigma_ch3

    if auto_t2 < TRUST_LIMIT_CH2:
        fixed_t2, t2_mode = FALLBACK_CH2, "FALLBACK"
    else:
        fixed_t2, t2_mode = auto_t2, "AUTO"

    if auto_t3 < TRUST_LIMIT_CH3:
        fixed_t3, t3_mode = FALLBACK_CH3, "FALLBACK"
    else:
        fixed_t3, t3_mode = auto_t3, "AUTO"

    lod_int_ch2 = fixed_t2 * min_cluster_size
    lod_int_ch3 = fixed_t3 * min_cluster_size
    lod_area_px = min_cluster_size

    print(f" > Threshold Ch2: {fixed_t2:.2f} ({t2_mode})")
    print(f" > Threshold Ch3: {fixed_t3:.2f} ({t3_mode})")

    # --- 3. PROCESSING ---
    per_cell_data = []
    global_image_data = []

    for condition_folder in sorted(base_path.iterdir()):
        if not condition_folder.is_dir() or condition_folder.name.startswith("Analysis_Run"): continue
        print(f"   --> FOLDER: {condition_folder.name}")

        for dapi_path in condition_folder.glob("*ch00.tif"):
            plt.close('all')
            img_dapi = io.imread(dapi_path, as_gray=True)
            img_ch2 = io.imread(str(dapi_path).replace("ch00.tif", "ch01.tif"), as_gray=True)
            img_ch3 = io.imread(str(dapi_path).replace("ch00.tif", "ch02.tif"), as_gray=True)

            thresh_d = filters.threshold_li(img_dapi)
            nuc_mask = morphology.remove_small_objects(img_dapi > thresh_d, 50)
            labels = measure.label(nuc_mask)
            dist = ndi.distance_transform_edt(~nuc_mask)
            cyto_labels = segmentation.watershed(-dist, labels, mask=dist < max_dist)
            cytosol_only = np.where(nuc_mask, 0, cyto_labels)

            mask_ch2 = morphology.remove_small_objects(img_ch2 > fixed_t2, min_size=min_cluster_size)
            mask_ch3 = morphology.remove_small_objects(img_ch3 > fixed_t3, min_size=min_cluster_size)

            total_cyto_mask = cytosol_only > 0
            total_cell_mask = total_cyto_mask | nuc_mask

            total_nuc_area = np.sum(nuc_mask)
            total_cyto_area = np.sum(total_cyto_mask)
            total_cell_area = np.sum(total_cell_mask)

            # Channel 2 Metrics
            i2_nuc, a2_nuc = np.sum(img_ch2[mask_ch2 & nuc_mask]), np.sum(mask_ch2 & nuc_mask)
            i2_cyto, a2_cyto = np.sum(img_ch2[mask_ch2 & total_cyto_mask]), np.sum(mask_ch2 & total_cyto_mask)
            i2_tot, a2_tot = i2_nuc + i2_cyto, a2_nuc + a2_cyto

            # Channel 3 Metrics
            i3_nuc, a3_nuc = np.sum(img_ch3[mask_ch3 & nuc_mask]), np.sum(mask_ch3 & nuc_mask)
            i3_cyto, a3_cyto = np.sum(img_ch3[mask_ch3 & total_cyto_mask]), np.sum(mask_ch3 & total_cyto_mask)
            i3_tot, a3_tot = i3_nuc + i3_cyto, a3_nuc + a3_cyto

            global_image_data.append({
                'Condition': condition_folder.name, 'Image': dapi_path.name,
                'Area_Cyto_px': total_cyto_area,
                # Ch2
                'Ch2_Cyto_Area_Fract': (a2_cyto/total_cyto_area*100) if total_cyto_area>0 else 0,

                # Ch3
                'Ch3_Cyto_Area_Fract': (a3_cyto/total_cyto_area*100) if total_cyto_area>0 else 0,
            })

            props = measure.regionprops(cyto_labels)
            for p in props:
                c_id = p.label
                m_n, m_c = (labels == c_id), (cytosol_only == c_id)
                m_tot = m_n | m_c
                per_cell_data.append({
                    'Condition': condition_folder.name, 'Image': dapi_path.name, 'Cell_ID': c_id,
                    'Ch3_Cyto_IntInt': np.sum(img_ch3[m_c & mask_ch3]), 'Ch3_Nuc_IntInt': np.sum(img_ch3[m_n & mask_ch3]), 'Ch3_Total_IntInt': np.sum(img_ch3[m_tot & mask_ch3])
                })

            # QC Outputs
            fig_h, ax_h = plt.subplots(1, 2, figsize=(10, 4))
            ax_h[0].hist(img_ch2[total_cell_mask].ravel(), bins=100, log=True, color='gray')
            ax_h[0].axvline(fixed_t2, color='r', linestyle='--'); ax_h[0].set_title(f"Ch2 ({t2_mode})")
            ax_h[1].hist(img_ch3[total_cell_mask].ravel(), bins=100, log=True, color='gray')
            ax_h[1].axvline(fixed_t3, color='r', linestyle='--'); ax_h[1].set_title(f"Ch3 ({t3_mode})")
            fig_h.savefig(hist_dir / f"Hist_{dapi_path.stem}.png"); plt.close(fig_h)

            fig_p, ax_p = plt.subplots(1, 3, figsize=(18, 6))
            ax_p[0].imshow(img_dapi, cmap='gray'); ax_p[0].contour(nuc_mask, colors='cyan', linewidths=0.5)
            ax_p[1].imshow(np.where(mask_ch2, img_ch2, 0), cmap='magma'); ax_p[1].contour(total_cyto_mask, colors='white', linewidths=0.3)
            ax_p[2].imshow(np.where(mask_ch3, img_ch3, 0), cmap='magma'); ax_p[2].contour(total_cyto_mask, colors='white', linewidths=0.3)
            for a in ax_p: a.axis('off')
            fig_p.savefig(preview_dir / f"QC_{dapi_path.stem}.png", bbox_inches='tight', dpi=150); plt.close(fig_p)
            gc.collect()

    # --- 4. EXPORT ---
    if global_image_data:
        df_cell, df_global = pd.DataFrame(per_cell_data), pd.DataFrame(global_image_data)
        df_filtered = df_global[~df_global['Image'].str.contains('2nd', case=False)]

        final_summary = pd.concat([
            df_filtered.groupby('Condition').mean(numeric_only=True).add_suffix('_mean'),
            df_filtered.groupby('Condition').std(numeric_only=True).add_suffix('_sd')
        ], axis=1).reset_index()

        avg_cyto_area = df_global['Area_Cyto_px'].mean()
        final_summary['LoD_AreaFract_Cyto'] = (lod_area_px / avg_cyto_area * 100) if avg_cyto_area > 0 else 0


        with pd.ExcelWriter(run_dir / "Final_Quantification_Results.xlsx") as writer:
            df_cell.to_excel(writer, sheet_name='Per_Cell_Data', index=False)
            df_global.to_excel(writer, sheet_name='Global_Image_Totals', index=False)
            final_summary.to_excel(writer, sheet_name='Condition_Summary', index=False)
            pd.DataFrame({'Parameter': ['Ch2_T', 'Ch3_T', 'Ch2_Mode', 'Ch3_Mode', 'MinSize', 'LoD_Int2', 'LoD_Int3'],
                         'Value': [fixed_t2, fixed_t3, t2_mode, t3_mode, min_cluster_size, lod_int_ch2, lod_int_ch3]}).to_excel(writer, sheet_name='Metadata', index=False)

        print(f"[{datetime.now().strftime('%H:%M:%S')}] Success! Analysis in: {run_dir.name}")
```

**Supplementary Code 2: Colocalization**

Description: The following script was used to calculate the Pearson Correlation Coefficient (*r*) between fluorescent channels

```
import numpy as np
import pandas as pd
from pathlib import Path
import matplotlib.pyplot as plt
from datetime import datetime
from skimage import io
from skimage.filters import threshold_otsu
from scipy.stats import pearsonr

"""
Script for calculating Pearson Correlation Coefficients in IF images.
Used for colocalization analysis in
'Influenza A virus NS1 sequesters RNA:DNA hybrids to evade RNase H1-dependent innate immunity'
"""

def compute_colocalization_from_channels(
    ch1_path, ch2_path, ch3_path,
    normalize=True,
    adaptive_threshold=None,   # 'otsu', 'percentile', or None
    fixed_threshold=None,      # used if adaptive_threshold is None
    return_pixels_above=False, # if True, return arrays of pixels above threshold
    verbose=False
):
    """
    Compute Pearson and Manders colocalization between three channels.

    Returns:
        results: dict of Pearson coefficients
        channels: list of np.arrays (normalized)
        thresholds: list of thresholds for each channel
        pixels_above: list of np.arrays with pixels above threshold (optional)
    """

    # --- Load images ---
    channels = []
    for i, path in enumerate([ch1_path, ch2_path, ch3_path]):
        ch = io.imread(path).astype(np.float32)
        # Convert RGB to grayscale if needed (assume single-channel in R, G, or B)
        if ch.ndim == 3:
            ch = ch.mean(axis=-1)
            if verbose:
                print(f"Channel {i+1} is RGB, converted to grayscale")
        # Normalize
        if normalize:
            ch_min, ch_max = ch.min(), ch.max()
            if ch_max > ch_min:
                ch = (ch - ch_min) / (ch_max - ch_min)
            else:
                ch = np.zeros_like(ch)
        channels.append(ch)

    ch1, ch2, ch3 = channels

    # --- Determine thresholds ---
    thresholds = []
    for ch in channels:
        if adaptive_threshold == 'percentile':
            t = np.percentile(ch, 95)
        elif adaptive_threshold == 'otsu':
            t = threshold_otsu(ch)
        else:  # fixed threshold
            t = fixed_threshold if fixed_threshold is not None else 0
        thresholds.append(t)

    t1, t2, t3 = thresholds

    if verbose:
        print("\n=== Thresholds per channel ===")
        for i, (ch, t) in enumerate(zip(channels, thresholds), start=1):
            print(f"Channel {i}: min={ch.min():.3f}, max={ch.max():.3f}, mean={ch.mean():.3f}, threshold={t:.3f}")

    # --- Flatten channels for Pearson ---
    flat1, flat2, flat3 = ch1.ravel(), ch2.ravel(), ch3.ravel()

    pearson_12, _ = pearsonr(flat1, flat2)
    pearson_13, _ = pearsonr(flat1, flat3)
    pearson_23, _ = pearsonr(flat2, flat3)

    results = {
        "Pearson_ch1ch2": pearson_12,
        "Pearson_ch1ch3": pearson_13,
        "Pearson_ch2ch3": pearson_23,
    }

    # --- Pixels above threshold ---
    pixels_above = None
    if return_pixels_above:
        pixels_above = [ch[ch > t] for ch, t in zip(channels, thresholds)]

    if return_pixels_above:
        return results, channels, thresholds, pixels_above
    else:
        return results, channels, thresholds

# --- Main Execution Block ---
input_folder = Path("path/to/your/images") #UPDATE THIS BEFORE RUNNING
datestamp =datetime.now().strftime("%Y%m%d")
output_csv = input_folder / f"{datestamp}_colocalization_results.csv" # Saved INSIDE input folder

preview_folder = input_folder / "previews"
preview_folder.mkdir(exist_ok=True)

all_results = []
anchor_files = sorted(input_folder.glob("*ch00.tif"))

print(f"Starting analysis of {len(anchor_files)} sets...")

for ch00_path in anchor_files:
    # 1. Determine paths
    base_str = str(ch00_path).replace("ch00.tif", "")
    ch01_path = Path(f"{base_str}ch01.tif")
    ch02_path = Path(f"{base_str}ch02.tif")

    if ch01_path.exists() and ch02_path.exists():
        file_id = ch00_path.stem.replace("ch00", "")

        # Simple print as requested
        print(f"Analyzing: {file_id}")

        # 2. Run function (verbose=False to hide the long logs)
        # We set return_pixels_above=True to get the pixel counts
        res, chans, threshs, pix_above = compute_colocalization_from_channels(
            ch00_path, ch01_path, ch02_path,
            adaptive_threshold='otsu',
            return_pixels_above=True,
            verbose=False
        )

        # 3. Create and Save Preview Image
        fig, axes = plt.subplots(2, 3, figsize=(15, 8))
        fig.suptitle(f"Verification: {file_id}", fontsize=16)
        titles = ["Ch0 (DAPI)", "Ch1", "Ch2"]

        for i in range(3):
            axes[0, i].imshow(chans[i], cmap='gray')
            axes[0, i].set_title(f"Original {titles[i]}")
            axes[0, i].axis('off')

            mask = chans[i] > threshs[i]
            axes[1, i].imshow(mask, cmap='magma')
            axes[1, i].set_title(f"Mask (Thresh: {threshs[i]:.3f})")
            axes[1, i].axis('off')

        plt.tight_layout()
        plt.savefig(preview_folder / f"{file_id}_preview.png")
        plt.close()

        # 4. Prepare data for CSV (including min, max, mean, threshold, and pixel count)
        row = {
            "Filename": file_id,
            # Correlation Metrics
            "Pearson_1-2": res["Pearson_ch1ch2"],
            "Pearson_1-3": res["Pearson_ch1ch3"],
            "Pearson_2-3": res["Pearson_ch2ch3"],
        }

        # Dynamically add Ch info (Min, Max, Mean, Thresh, PixelsAbove)
        for i in range(3):
            ch_num = i
            row[f"Ch{ch_num}_Min"] = chans[i].min()
            row[f"Ch{ch_num}_Max"] = chans[i].max()
            row[f"Ch{ch_num}_Mean"] = chans[i].mean()
            row[f"Ch{ch_num}_Threshold"] = threshs[i]
            row[f"Ch{ch_num}_PixelsAbove"] = pix_above[i].size

        all_results.append(row)

# 5. Finalize CSV
if all_results:
    pd.DataFrame(all_results).to_csv(output_csv, index=False)
    print(f"\nSuccess! Results saved to: {output_csv}")
```
