## Supplementary figures and images for "Influenza A virus NS1 sequesters RNA:DNA hybrids to evade RNase H1-dependent innate immunity"

### Supplementary Figures

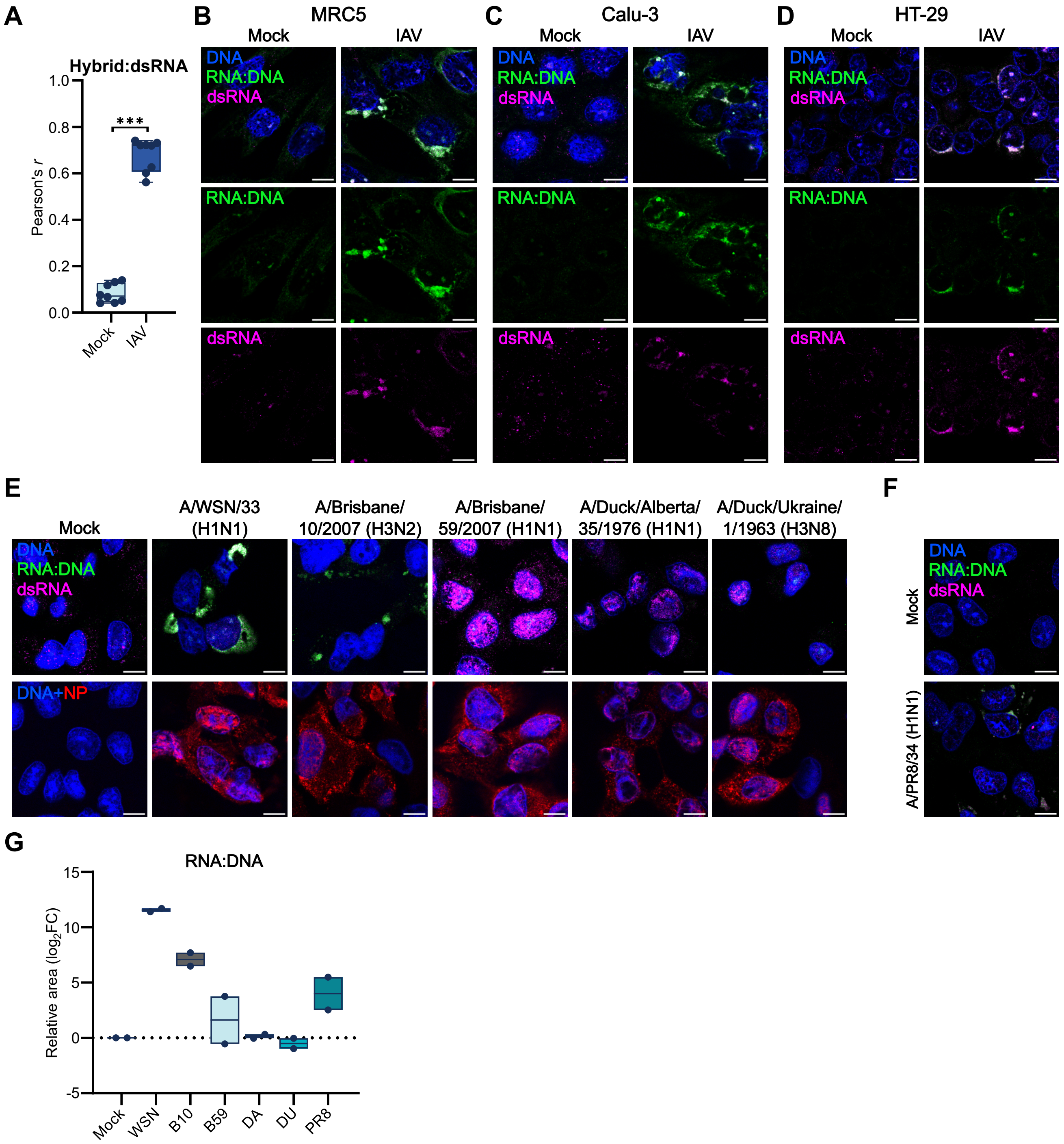

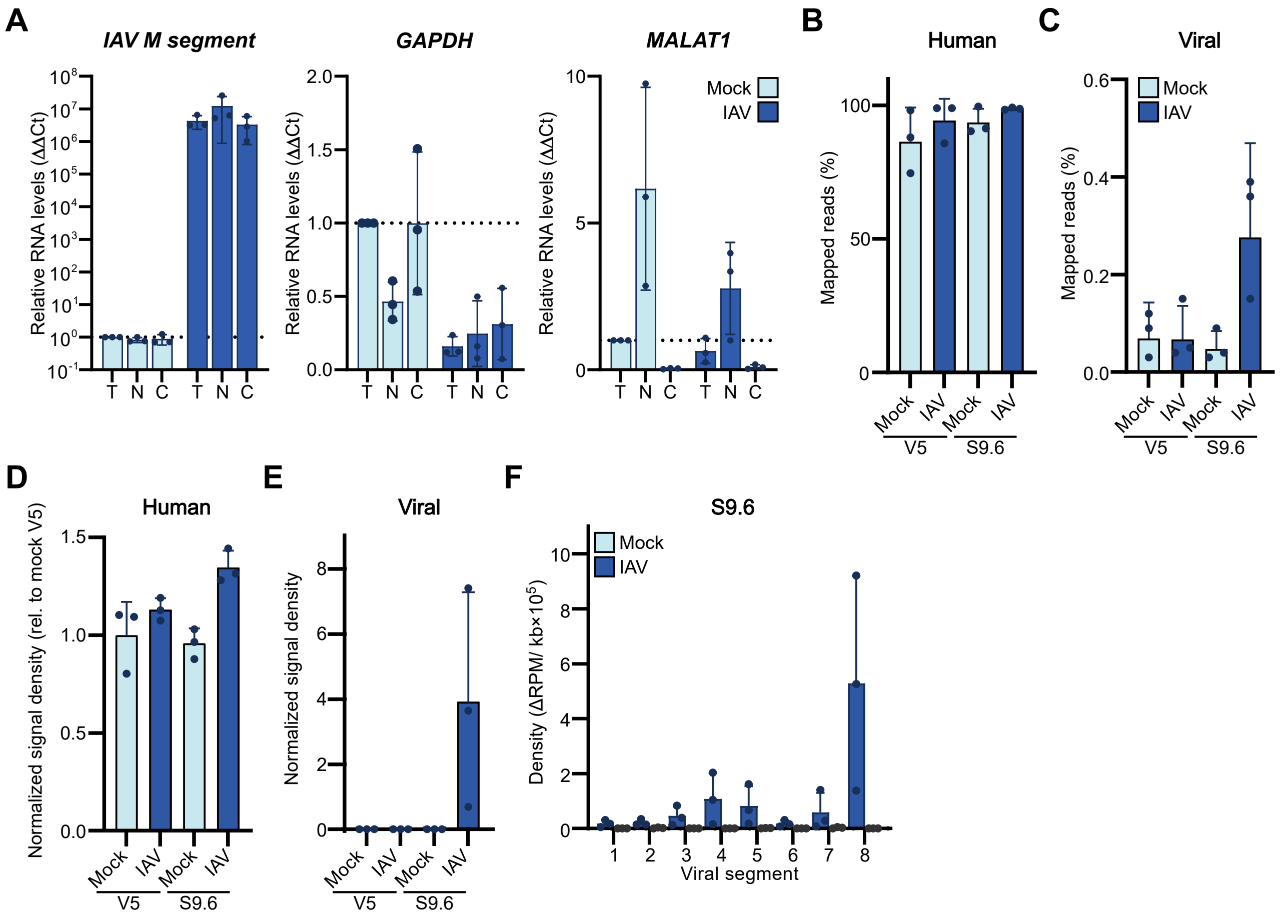

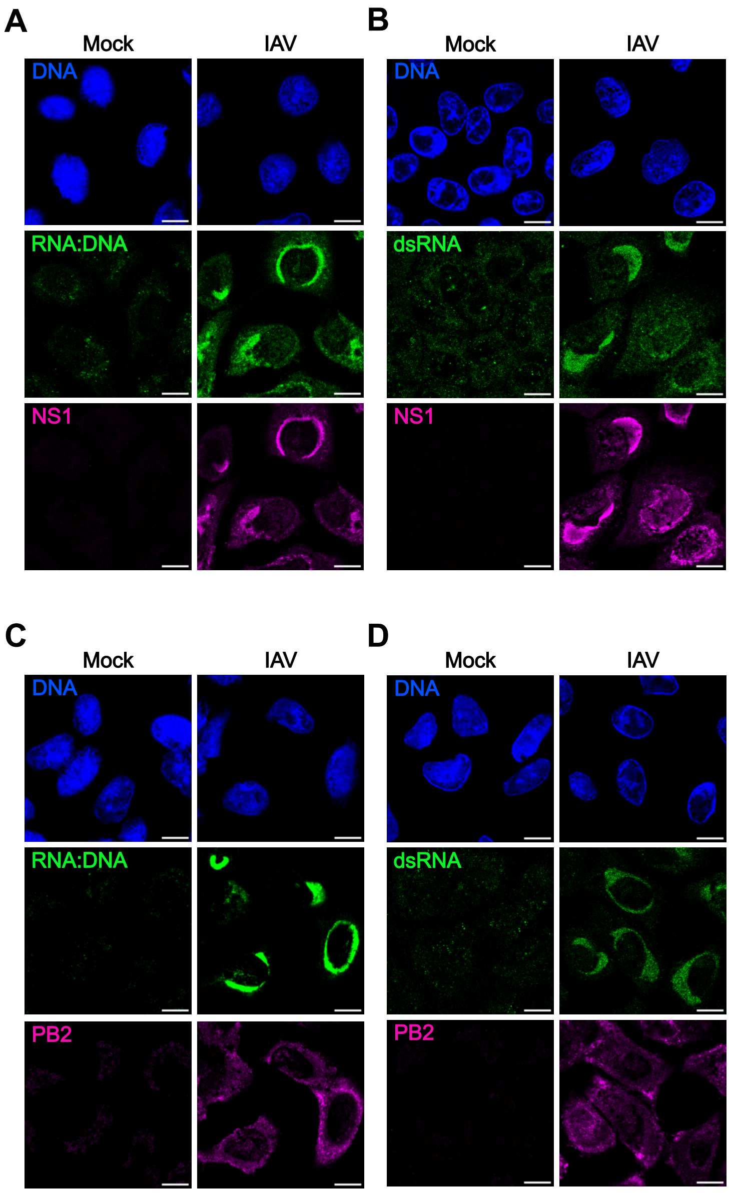

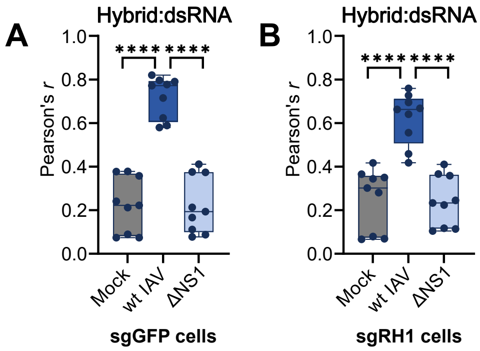
